# Direction mutation pressure of SARS-CoV-2 helps to understand the past and predict the future evolution: C>U and G>U biased mutagenesis forces the majority of amino-acid substitutions to be from CG-rich losers to U-rich gainers

**DOI:** 10.1101/2022.08.22.504819

**Authors:** Alexandr Voronka, Bogdan Efimenko, Sergey Oreshkov, Melissa Franco, Zoe Fleischmann, Valerian Yurov, Arina Trufanova, Valeria Timonina, Natalia Ree, Arthur Zalevsky, Emma Penfrat, Thomas Junier, Alexey Agranovsky, Konstantin Khrapko, Konstantin Gunbin, Jacques Fellay, Konstantin Popadin

## Abstract

Evolution is a function of mutagenesis and selection. To analyse the role of mutagenesis on the structure of the SARS-CoV-2 genome, we reconstructed the mutational spectrum, which was highly C>U and G>U biased. This bias forces the SARS-CoV-2 genome to become increasingly U-rich unless selection cancels it. We analysed the consequences of this bias on the composition of the most neutral (four-fold degenerate synonymous substitutions) and the least neutral positions (nonsynonymous substitutions). The neutral nucleotide composition is already highly saturated by U and, according to our model, it is at equilibrium, suggesting that in the future, we don’t expect any more increase in U. However, nonsynonymous changes continue slowly evolve towards equilibrium substituting CG-rich amino-acids (“losers”) with U-rich ones (“gainers”). This process is universal for all genes of SARS-CoV-2 as well as for other coronaviridae species. In line with the direction mutation pressure hypothesis, we show that viral-specific amino acid content is associated with the viral-specific mutational spectrum due to the accumulation of effectively neutral slightly deleterious variants (losers to gainers) during the molecular evolution. The tuning of a protein space by the mutational process is expected to be typical for species with relaxed purifying selection, suggesting that the purging of slightly-deleterious variants in the SARS-CoV-2 population is not very effective, probably due to the fast expansion of the viral population during the pandemic. Understanding the mutational process can help to design more robust vaccines, based on gainer-rich motifs, close to the mutation-selection equilibrium.

## INTRODUCTION

Molecular evolution is a function of mutagenesis and selection. Deconvolution of these two factors is an essential step for a deep understanding of the whole process. Mutagenesis is far from random: a set of tissue and species-specific mutagens determines highly nonuniform probabilities of nucleotide substitutions. The mutational spectrum, defined here as a relative rate of single nucleotide substitutions in the neutral region of nucleic acid, allows characterising and comparing the mutational processes. Depending on the depth of input data, the mutational spectrum can be reconstructed with various precision: 2 components, the ratio of transitions to transversions (1, 2); 6 components, single nucleotide substitutions with symmetrical mutagenesis on both strands (where, for example, C>T is equivalent to G>A) (3); 12 components, single nucleotide substitutions with strand-specific mutagenesis (where C>T is not equal to G>A) (4); 96 components, the most commonly accepted spectrum in the cancer field, assumes symmetrical single nucleotide substitutions with trinucleotide context (4×6×4=96) (5, 6); 192 components, strand-specific single nucleotide substitutions with trinucleotide context (4*12*4) (7, 8)or even more.

The mutational spectrum, after reconstruction, can be used for various purposes. The most important implication is tracing potential mutagens through their mutational signatures (9). The mutational spectrum, interpreted as a new molecular phenotype of cancer patients, can have several clinical applications soon (10). Although historically, the progress in the analysis of the deep mutational spectra was associated with human cancers, currently, it goes beyond it and expands to all species. For example, several recent studies of the SARS-CoV-2 mutational spectrum allowed to understand its evolution better: high similarities between the mutation spectra of SARS-CoV-2 and the bat coronavirus RaTG13, suggested the natural origin of SARS-CoV-2 (11); asymmetry and directionality of the mutational spectrum observed in SARS-CoV-2 enabled authors to trace the root of the tree (8); the mutational spectrum helped better to infer selection and suggest a way to design more robust vaccines (12).

Long-term consequences of the mutational spectrum lead to several fundamental questions: (i) how strongly do mutational spectra affect the nucleotide composition in neutral (for example, in synonymous positions) and not neutral (for example, nonsynonymous) sites; (ii) how far different species (genomes and genic regions) are from their mutation-selection equilibrium; (iii) how often mutational spectra is changing along the phylogeny? The reconstruction of the mitochondrial (mtDNA) mutational spectrum of hundreds of vertebrate species allowed to address some of these questions and demonstrated that synonymous positions of mammalian mtDNA (4)and mtDNA of Actinopterygii (13)are strongly affected by the mutational spectrum and close enough to equilibrium. One of the most famous and still unsolved discrepancies between the mutational spectrum and the genome-wide nucleotide content is the increased GC content of aerobic versus anaerobic bacteria (14–16): according to the well-known hallmark of oxidative damage, G>T is expected to be higher in aerobic versus anaerobic bacteria, making genomes of the former more GC-poor and AT-rich; however the observed trend is opposite: aerobic bacteria have a higher GC content as compared to anaerobic (14–16). It is important to emphasise also that mutational bias can change nucleotide composition not only in neutral sites but also in nonneutral ones - affecting the amino-acid composition of species (17–19).

Here to analyse the consequences of the mutational spectrum on neutral and non-neutral sites of a genome, we focused on SARS-CoV-2, which is advantageous because of: (i) strongly asymmetric mutational spectrum; (ii) high absolute mutational rate, typical for all RNA viruses; (iii) an expansion phase during the pandemic, which is known to be associated with inefficient selection and accumulation of slightly-deleterious substitutions (20, 21); (iv) large scale dataset, allowing to reconstruct detailed mutational spectrum (22). First, we confirmed the highly biased towards U mutational spectrum; we demonstrated that synonymous positions are very close to the compositional equilibrium and nonuniformity in codon usage is shaped by the mutational spectrum. Second, we observed an effect of the mutational spectrum on the most common amino acid substitutions, which predominantly happen from CG-rich amino acids (“losers”) to U-rich amino acids (“gainers”). Finally, analysing the amino-acid composition of all single-stranded positive RNA viruses, we observed that all coronaviridae have an excess of gainers versus losers, assuming that the coronavirus-specific mutational spectrum shapes its amino-acid content. We propose the butterfly effect scenario when little changes in the mutational spectrum (which presumably occurred in ancestral coronaviridae) in a very long-term perspective lead to drastic changes in the amino-acid composition. We hypothesise that the butterfly effect can be the strongest in species with the asymmetric mutational spectrum, high absolute mutation rate and relaxed purifying selection. We believe that the progress in understanding the permissive (weekly selected) amino-acid trajectories (23)will help deconvolute the mutational bias’s effect on amino-acid composition.

## MATERIALS AND METHODS

### Data collection and processing

A set of 4,339,984 full-length SARS-CoV-2 genome sequences was downloaded on 14.10.21 from GISAID (https://www.epicov.org). Duplicates and low-quality sequences containing uncertain nucleotides (N, K, H etc.) or less than 29001 nucleotides were removed. Filtered 1,139,387 sequences were aligned to Wuhan-Hu-1 reference genome (NC_045512.2) with bwa bwasw command (v.0.7.17-r1188) (24). Alignment was used for phylogenetic tree building and different descriptive statistics. The phylogenetic tree was constructed using FastTree 2.1.10 (25) on a sample set of aligned 203,045 genomes. The complete tree was pruned to 54,521 nodes with a bottom-up approach to conserve more phylogenetic differences in the tree. Briefly, pruning consists of iterative deleting of nodes in terminal polytomies and random terminal nodes until the number of tree nodes is less than 55000. Ancestral genomes of pruned tree internal nodes were reconstructed using PRANK v.170427 (26) using the default model of DNA sequence evolution (HKY model). PRANK uses BppAncestor library (27)for the fast marginal maximum likelihood reconstruction of the most likely ancestral sequence at each node. This type of maximum likelihood ancestral reconstruction goes from the descendants progressively assigning the most likely character state to each ancestor based on only its immediate descendants and considering the user-defined sequence evolution model. This approach does not consider the ancestral states’ probability joint distribution in each alignment position and tree node. Therefore, it often converges to a local optimum (28), leading to overestimating the substitution number. However, this approach is fast enough for feasible computations of alignment consisting of several tens of thousands of viral genomes.

Single nucleotide substitutions from each node in the tree were called taking into account that upstream and downstream two-nucleotide neighbour positions were the same and did not contain deletions.

### Mutational spectrum reconstruction

The mutation spectrum of SARS-CoV-2 was based on a sample of substitutions collected from the phylogenetic tree. We calculated mutational profiles of the dataset of mutations, i.e. counted all possible variants of single nucleotide substitutions (12 numbers or 192 numbers in case of using the neighbouring nucleotides) and adjusted them to the expected numbers of substitutions that were collected from the reference sequence Wuhan-Hu-1. When a mutation sample was collected from a distinct genome region or was based, for example, on synonymous substitutions, we adjusted the calculated mutational profile to corresponding expected substitutions numbers.

### Comparison of mutation ratios in mutational spectra

Pairs of normalised values of 12-component spectra were compared using a binomial test to test imbalances in mutational spectra (29). We compared only mutually exclusive subtitutions, i.e. reciprocal pairs (C>U and U>C etc.) and complementary pairs (C>U and G>A etc.); consequently the number of mutations in such pairs might be considered as binomially distributed. P-values were corrected with the Benjamini and Hochberg procedure.

### A sampling of late and early genomes

From the entire dataset of genome sequences downloaded from GISAID, we extracted three groups of sequences split by date: from December 2019 to March 2020 (early), from the beginning of October 2020 to the end of October 2020 (intermediate) and from September 2021 to October 2021 (late). From each group, we sampled randomly 10000 sequences and used them in the analyses to compare the nucleotide frequencies.

### Simulation of the nucleotide content

An expected neutral nucleotide composition, evolved under a 12-component mutational spectrum, was obtained using computational simulations and differential equations. Both approaches gave very similar results. Details are available in our recent manuscript (4).

### Analysis of codon usage of human single-stranded positive RNA viruses

From the NCBI virus database (https://www.ncbi.nlm.nih.gov/labs/virus/vssi/#/) we extracted RefSeqID of all human, completely sequenced single-stranded positive RNA viruses (N = 149 - segmented viruses are counted with each separate segment), downloaded all viral RefSeqs from ftp ncbi (ftp://ftp.ncbi.nlm.nih.gov/refseq/release/viral/) and estimated their genome-wide codon usage.

### Analysis of amino acid substitutions

To analyse how changes in amino acid composition depend on the mutation spectrum, we used nonsynonymous substitutions. We separated all amino acid substitutions by mutational spectrum: direct substitutions (caused by the main substitutions C>U and G>U) and opposite substitutions (caused by U>C and U>G). Then to count whether the amino acid composition changes, we took the number of direct substitutions, for example, Val>Phe (GUU>UUU or GUC>UUC), and the number of opposite substitutions, Phe>Val (UUU>GUU or UUC>GUC). Using these numbers, we calculated how much the fractions of direct substitutions differ from the random value (0.5) using a binomial test.

### RdRp dataset

From the Serratus virus database (30), Trees and Alignments section (https://serratus.io/trees), we downloaded alignments of RNA-dependent RNA polymerase sequences of (+)ssRNA viruses. We estimated amino acid usage for each sequence and performed PCA on the obtained dataset.

### Data access

Data and scripts are available on our GitHub: https://github.com/mitoclub/SARS-CoV-2-MutSpec.

## RESULTS

### 1. The mutational spectrum of SARS-CoV-2 demonstrates strong strand-asymmetry and directionality both leading to an increase in U

To uncover the main patterns of molecular evolution of SARS-CoV-2 it is important to distinguish mutagenesis from a selection. To do it, we reconstructed the mutational spectrum of SARS-CoV-2 by the following pipeline (see methods for details):

i. sampled 1,139,387 whole-genome sequences out of 4,339,984 available at GISAID at 14.10.2021;
ii. aligned genomes against the Wuhan-Hu-1 reference genome and pruned the set to 54,521 sequences to make the following steps computationally feasible;
iii. reconstructed phylogenetic tree and ancestral sequences in each internal node;
iv. counted all observed single nucleotide substitutions and normalised them on nucleotide content (for each type of substitution, we divided the number of observed mutations by the number of expected ones). This normalisation step eliminates the effect of different nucleotide compositions, which helps uncover asymmetry and directionality and makes the mutational spectra comparable between different organisms and signatures.

As a result of our pipeline, we got 542 768 high-quality (see methods) single-nucleotide substitutions, among which 314 538 were nonsynonymous, 206 319 synonymous and 92734 synonymous at four-fold positions. Next, we reconstructed three types of the mutational spectrum of SARS-CoV-2: (i) “ALL” is a spectrum based on all mutations, which assures the largest sample size with the cost of potential selection signal; (ii) “SYN” is a spectrum based on synonymous mutations only, which is more neutral but less numerous and (iii) “SYN4F” is a spectrum based on synonymous mutations in four-fold degenerate sites which is the most neutral subset of mutations but, in parallel, is the least numerous one (some contexts are missing due to the structure of the genetic code).

All three mutational spectra showed (i) substantial excess of C>U substitutions, (ii) strand-asymmetry and (iii) directionality. C>U substitutions contributed the majority of substitutions: 37% for ALL (Fig 1A), 25% for SYN and 35% for SYN4F (Fig. S1a-b). The mutational spectrum was strand-asymmetrical in terms of the complementary base pairing: if a probability of a given nucleotide mutating into another one (for example C>U) is the same on all strands, frequencies of C>U on the positive strand would be similar to C>U on the negative strand, which is equivalent to G>A on the positive strand. However, the observed frequencies of such complementary substitutions on positive strand were different with a strong excess of C>U as compared to G>A and G>U as compared to C>T (Fig 1A, Fig. S1a-b), suggesting that there is either a strand difference in mutagenesis or reparation. The mutational spectrum was also directional in terms of the rates of reciprocal mutation, i.e. mutated and substituted bases: for example, C>U is higher than U>C, and G>U is more elevated than U>G (ALL - Fig 1A; SYN and SYN4F - Fig. S1a-b; Supplementary Tables 2-4, results of the binomial test) suggesting that, in the absence of selection, an increase in U is expected.

**Fig. 1.**
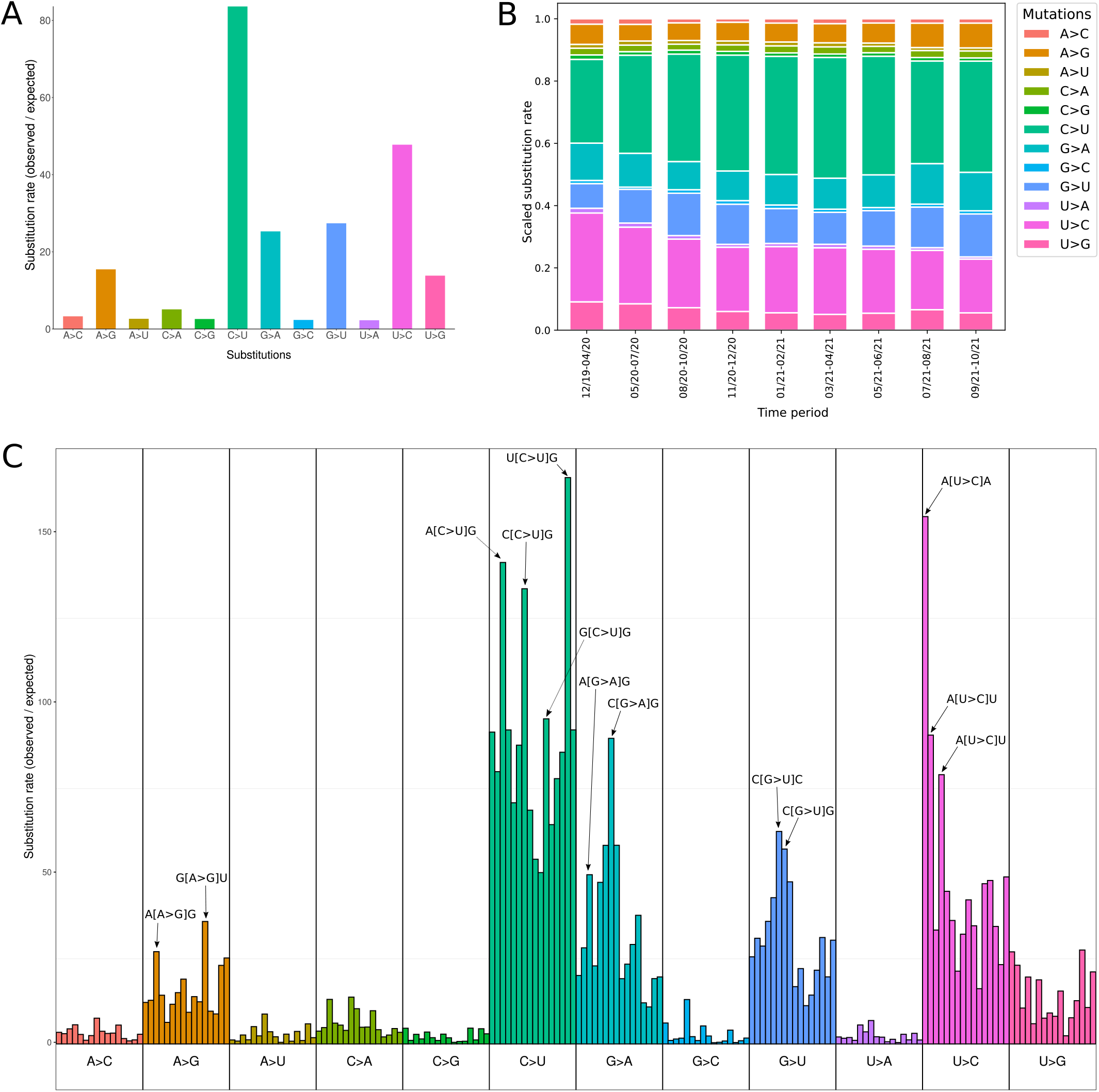
Mutational spectrum of SARS-CoV-2 based on an ALL set of mutations. All substitutions are presented in the positive strand notation, which is a reference sequence of the SARS-CoV-2. The colour code (12 types of substitutions) is the same in all panels. 1A The 12-component mutational spectrum of SARS-CoV-2. For the figure, we used the set of mutations ALL. 1B Changes in the 12-component mutational spectrum of SARS-CoV-2 over pandemic. For the figure, we used the set of mutations ALL. 1C 192-component mutational spectrum of SARS-CoV-2 reconstructed for all nucleotides. For the figure, we used the set of mutations ALL.

To check the stability of the mutational spectra along the viral genome we reconstructed the mutational spectrum for each separate gene. We observed that the mutational spectrum was similar along the viral genome (Fig. S1c), suggesting that the key mutational process is not gene-specific but rather universal for the whole genome.

To check the stability of the mutational spectrum along the time of the pandemic, we reconstructed mutational spectra from nine time intervals of several months. To do it, we focused on genomes sequenced within those time intervals and used only terminal branches of the phylogenetic tree for the spectra reconstruction (see methods). We observed that (Fig 1B) ALL mutational spectrum changed through the pandemic: it became even more directional (C>U increased while U>C decreased; G>U increased while U>G decreased) and even more strand-asymmetrical (C>U increased while G>A didn’t change; G>U increased while C>A didn’t change). A similar pattern was also observed for SYN and SYN4F spectra (Fig. S1d-e). This finding couldn’t be explained by the graduate and low changes in the nucleotide composition of the viral genome and thus, it requires a special explanation, based either on changes in error-associated techniques, viral mutagenic properties or host mutagenic environment.

To evaluate the effect of the nucleotide context, which typically has a substantial impact on mutagenesis (10), we analysed the three-nucleotide context, resulting in the 192-component spectrum (4 neighbour nucleotides at 5’ x 12 types of substitutions x 4 neighbour nucleotides at 3’). ALL spectrum (Fig 1C) showed that C>U occurred preferentially in the context of [A/C/U]CG, while G>U substitution took place in the context [G/U]GU and GGA. Analysis of the 192-component spectra, reconstructed for SYN and SYN4F sets, demonstrated similar mutagenic motives (Fig. S1f-g).

### 2. The mutagenesis shapes the SARS-CoV-2 nucleotide content

Both strand-asymmetry and directionality of the SARS-CoV-2 mutational spectra (Fig 1, Fig. S2) are expected to increase the frequency of U unless the selection is strong enough to purge this effect. Do we observe an increase in U content during the pandemic or the mutational increase in U is completely erased by a selection, decreasing the frequency of U? To evaluate the effect of the mutational bias on the integral molecular evolution of the viral genome, we analyzed the changes in the nucleotide composition of the complete genome along the time of the pandemic. We randomly sampled genomes from three different time points (see fig 2A) and used a reference genome as a proxy for the most ancestral state. In line with our expectations, we observed an accumulation of U (an increase in 10 nucleotides between the first and the last time point), and a reduction of all other nucleotides (A, G and C) on the scale of the complete viral genome (dataset ALL, Fig 2A). It is important to emphasise that this integral analysis - a comparison of the whole-genome nucleotide content of random genomes, sequenced at certain time intervals - depended on a full mix of evolutionary forces, such as mutagenesis, selection and drift. Despite this, we observed that the changes in the whole-genome nucleotide composition follow the mutational bias, suggesting that mutagenesis is one of the strongest determinants of the current molecular evolution of SARS-CoV-2.

**Fig. 2.**
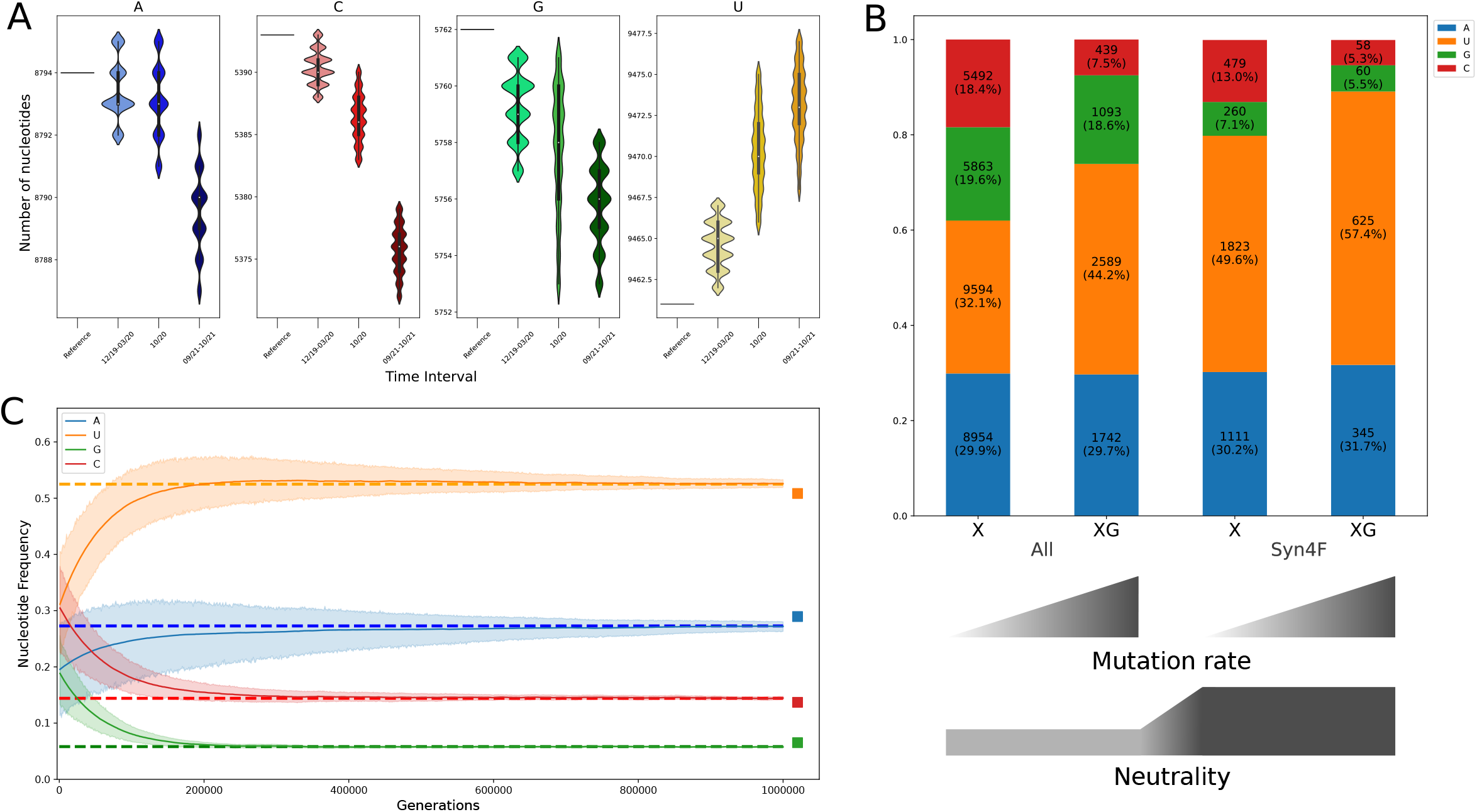
Changes in the nucleotide composition of SARS-CoV-2 over pandemic. The colour code (4 types of nucleotides) is the same in all panels. 2A Changes in the nucleotide composition of SARS-CoV-2 in all sites of the genome during the pandemic. The most left horizontal interval corresponds to the nucleotide composition of the reference sequence. 2B Nucleotide composition in the reference sequence of SARS-CoV-2. An excess of U is more prominent in sites with low selective constraints (SYN4F versus ALL) and a high substitution rate (G versus X). 2C Observed and expected nucleotide compositions are very similar. SYN4F dataset. Coloured shadows reflect changes in the nucleotide frequencies with the simulation time: they start at random positions and converge to their equilibrium state. Dash lines reflect the nucleotide composition of SYN4F in the reference sequence. Squares reflect an exact equilibrium state derived by the analytical approach with differential equations.

Interestingly, analyses of nucleotide composition based on the SYN4F set demonstrated no trend during the pandemic (Fig. S2a). This might be due to either the low number of such sites and weak statistical signal or the proximity of these sites to the equilibrium (i.e. saturation effect). To test the potential proximity of the sites to the nucleotide compositional equilibrium we compared four categories of sites characterized by different selective constraints and mutation rate: ALL NpN - all sites in the genome irrespectively of the nucleotide context (N is any nucleotide); ALL NpG - all sites in the genome preceding G (which is the hot spot for mutations C>U, see fig 1C); SYN4F NpN - four-fold degenerated synonymous sites irrespectively of the nucleotide context; SYN4F NpG - four-fold degenerated synonymous sites in the genome, preceding G. We observed a significant excess of U fraction in the least constrained (SYN4F versus ALL) and the most mutagenic sites (NpG versus NpN context, where N is any nucleotide) (Fig. 2B, Fig. S2b-c), suggesting that the nucleotide composition of the neutral and highly mutated sites is the closest to the expected equilibrium, which can be explained by the long-term evolution of a SARS-CoV-2 (even in previous hosts) under similar mutational pressure.

To estimate quantitatively the expected neutral (if we do not take into account the selection) nucleotide composition of the SARS-CoV-2, we performed a computer simulation based on a 12-component mutational spectrum of synonymous substitutions SYN4F and random initial nucleotide compositions (see methods). Interestingly, we observed that an equilibrium solution precisely reflects the observed nucleotide composition at SYN4F sites, (Fig 2C), confirming that SYN4F sites in SARS-CoV-2 genome has already been affected by the very similar mutational spectrum and this process was strong enough to shape the nucleotide composition towards the equilibrium. The analytical exact solution of the nucleotide compositional equilibrium, obtained with an input 12-component mutational spectrum, was completely in line with the simulation data and confirmed that SYN4F nucleotide composition is very close to equilibrium.

### 3. The codon usage of SARS-CoV-2 is affected by the mutational bias

The mutational bias affects the global nucleotide content of synonymous sites (Fig 2), which can be traced to the level of codon usage for individual amino acids. We plotted a standard genetic code as a heatmap with colour-coded numbers of codons from all protein-coding genes of the SARS-CoV-2 reference genome (Fig 3A). On the heatmap there are red horizontal stripes emphasising an increased number of NNU codons compared to NNC, an increased number of NNU codons compared to NNG and an increased number of NNA codons as compared to NNG. All these patterns can be explained by two of the most common asymmetrical mutations: C>U and G>U. The strongly asymmetrical codon usage of the reference genome, which is in line with the effect of the described mutational spectrum (Fig 1) provides additional lines of evidence that SARS-CoV-2 before the entering of the human host has evolved under a very similar mutational spectrum, suggesting that the key mutagens of SARS-CoV-2 are not host-specific, but rather viral-specific.

**Fig. 3.**
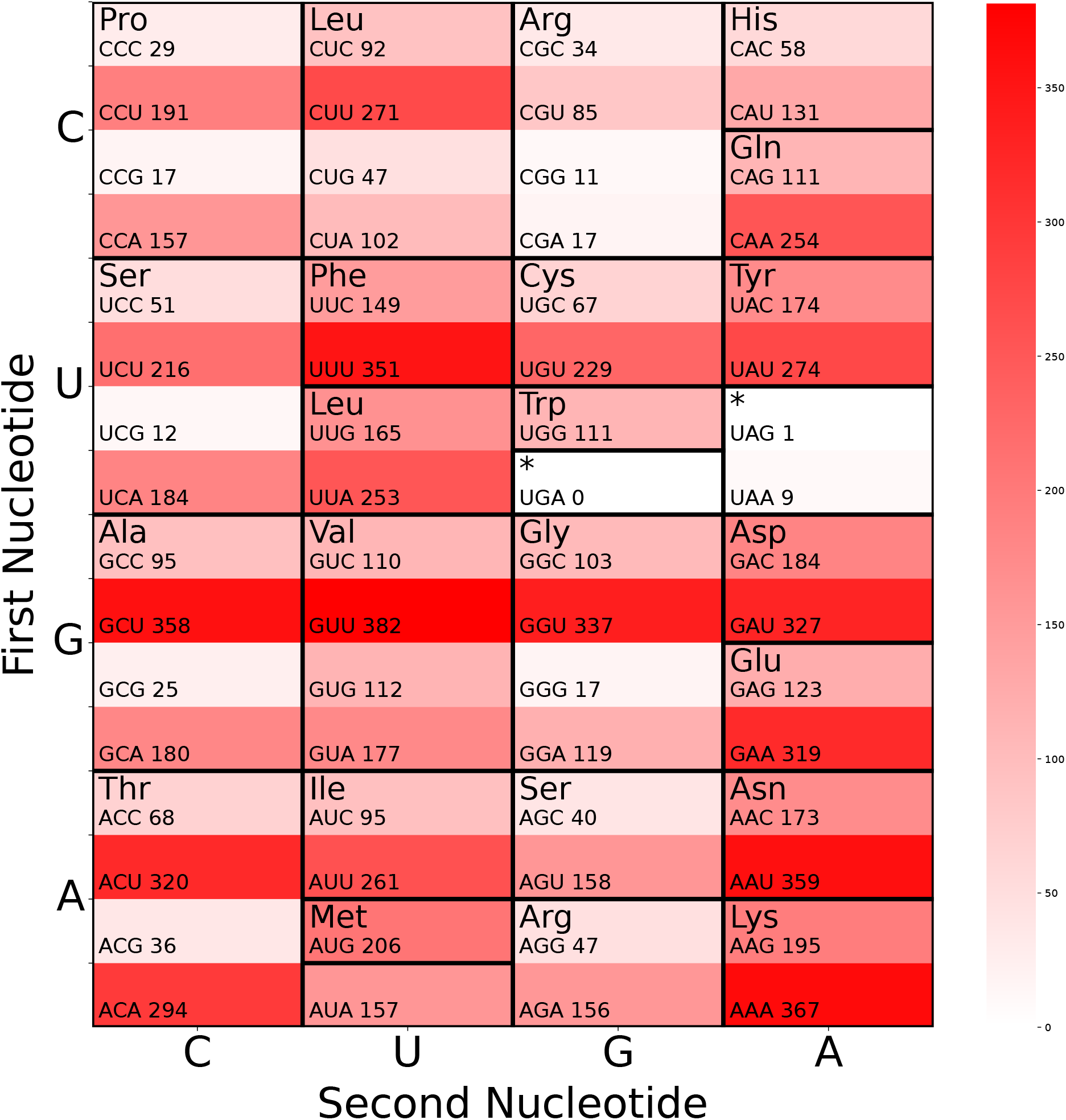
The codon usage of SARS-CoV-2 is affected by the mutational bias. 3A A heatmap of the codon usage in the SARS-CoV-2 reference genome. All codons of all protein-coding genes of the SARS-CoV-2 reference sequence were plotted as a heatmap. Horizontal stripes (increased frequency of NNU and NNA) can be explained by the mutational bias C>U, G>U and probably G>A.

Changes in the codon usage between early and late genomes during the pandemic did not show any significant trends (Fig. S3), which is in line with the equilibrium state in neutral positions (Fig 2C).

### 4. Amino acid substitutions are affected by the mutational bias

If selection is weak and the mutational spectrum is asymmetrical or directional, it may affect the pattern of amino acid substitutions. To test this, on the standard genetic code table, we plotted the expected trajectories of amino acid substitutions taking into account two of the most frequent asymmetrical substitutions: C>U and G>U (see the “butterfly scheme” in Fig 4A, named so due to a similarity with a butterfly). According to these trajectories, we categorised all amino acids into four groups: losers (Pro, Ala, Arg, Gly, Thr, Ser (AGX), Arg, His, Gln, Asp, Glu), gainers (Phe, Leu (UUX), Ile, Met, Tyr), intermediate (which are in the middle of the trajectory: Leu (CUX), Cys, Trp, Ser, Val) and neutral to the mutational bias (Asn, Lys). Next, we empirically tested these predictions using two types of data: (i) all amino acid substitutions reconstructed from the phylogenetic tree, (ii) changes in the amino acid composition between the late and early genomes.

**Fig. 4.**
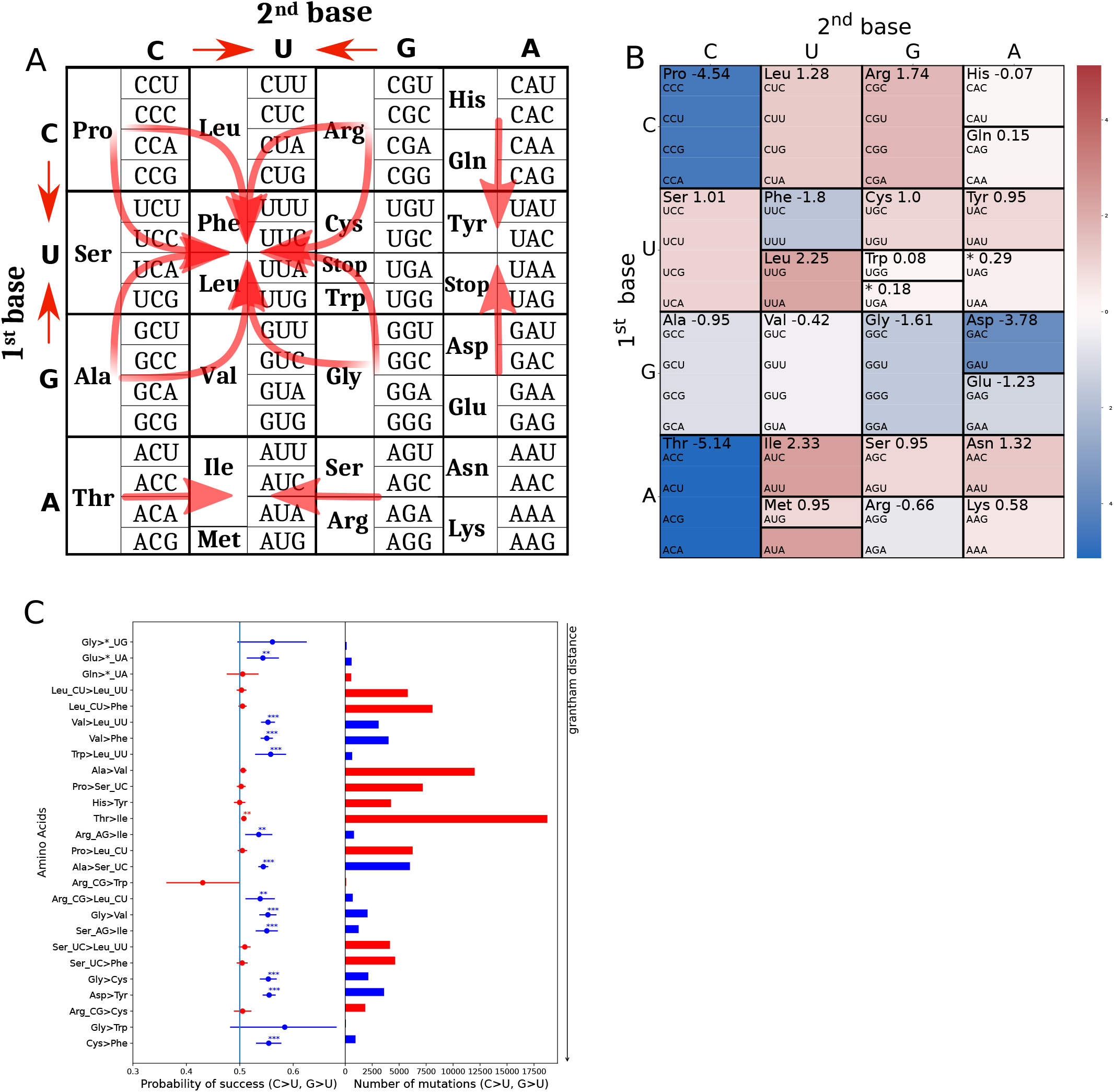
Amino acid substitutions of SARS-CoV-2 are affected by the mutational bias: a butterfly effect. 4A A “butterfly scheme” - expected amino acid trajectories driven by the C>U and G>U mutations in the space of the standard genetic code (indicated by red arrows). Arrows representing amino acid trajectories are drawn from the expected losers (Pro, Ala, Arg, Gly, Thr, Ser (AGX), His, Gln, Asp, Glu) to the expected gainers (Phe, Leu (UUX), Ile, Met, Tyr). 4B Average changes in amino acid composition over the course of the pandemic: increase (in red) in the frequency of gainers and decrease (in blue) in the frequency of losers in the late versus earlier genomes (see methods). 4C The pattern of the amino acid substitutions reconstructed from the phylogenetic tree. All substitutions are ranked correspondingly to the Grantham distance: from the amino acids closest to each other in physico-chemical terms, at the top, to the most distant ones at the bottom. Red colour marks substitutions shaped by C>U mutations, and blue - mutations shaped by G>U mutations. *_UG and *_UA denote stop codons UGA and UAA/UAG correspondingly. Asterisks mark statistically significant deviations from 0.5: ** p value < 0.01, *** p-value < 0.001 (binomial test, see methods and main text; all p-values are nominal).

Among all amino acid substitutions reconstructed on the phylogenetic tree (314538 mutations), the majority (193102 mutations) were located along the hot mutational trajectories (forward or backwards to the red arrows in Fig 4A). Among them, mutations concordant with the directionality of the mutagenesis (99689 mutations, ∼ 32% from all amino-acid substitutions, from losers to gainers or from losers to intermediate or from intermediate to gainers) were in 6.7% excess as compared to mutations, discordant with the direction of the mutational trajectories (93413 mutations from gainers to losers or from gainers to intermediate or from intermediate to losers). The difference in the directionality was statistically significant (p = 5.1e-9, binomial test). Interestingly, the directional effect was also significant for most individual pairs of amino acid substitutions (Fig 4C). For example, we observed 4019 mutations from Val to Phe and only 3280 from Phe to Val (Supplementary Table 5). Moreover, the same trend is visible not only at the level of all protein-coding genes but also separately within the ORF1AB region and structural protein-coding regions (Fig. S4B-C). An additional observation is that amino-acid substitutions shaped by C>U mutations (red colour in Fig 4C) are less biased as compared to substitutions, shaped by G>U mutations (blue colour in Fig 4C), which can be explained by the saturation effect: the majority of effectively-neutral amino-acid substitutions driven by the most common C>U substitutions already happened and the de novo ones are under purifying selection, while substitutions driven by G>U are far from saturation and still are accumulating. For example, Arg(CGX) shows no significant trend toward Cys and Trp (Fig 4C), probably because it is already close to the minimum frequency due to the mutational hot spot: CpG context is one of the strongest contexts to mutate towards UpG (Fig 1C).

After the origin of a mutation, its fate depends on the reproductive success of a carrier. In our previous “phylogeny tree” analyses (Fig 4C), we didn’t consider the frequency of descendants of each mutation, thus diminishing a potential effect of selection. Here, to trace the effect of the mutational bias on the global evolution of SARS-CoV-2, we run an integral analysis by sampling late and early genomes (see methods) and comparing their amino acid composition (Fig 4B). We observed that Pro, Thr and Asp demonstrated the most substantial decrease during the pandemic, while Leu (UUX) and Ile demonstrated the strongest increase in their numbers (Fig 4B). Interestingly, the dynamics of these five amino acids are consistent with the mutational trajectories: those with decreasing trends are losers, and those with increasing trends are gainers. (Fig 4A). Moreover, if we take into account eleven expected losers (Fig 4A) we observe that eight of them (all except Arg(CGN), Ser(AGN) and Gln) indeed increase in frequency at least moderately. Similarly, five of the six expected gainers show an increase (all except Phe). Altogether, we observe that even at the amino acid level, asymmetrical mutagenesis shapes the pattern of amino-acid changes between early and late SARS-CoV-2 genomes.

The recent and highly transmissible line of SARS-CoV-2, omicron, is characterised by the numerous nucleotide changes from the Wuhan reference sequence. To test whether the amino-acid substitutions associated with the evolution of omicron are in line with our hypothesis, we analysed all amino acid changes between the Wuhan and omicron sequences. Interestingly, among all amino acid changes (Supplementary Table 6), we observed that 12 amino acid substitutions were in line with the mutational bias. In contrast, only 3 - in the opposite direction (binomial test, the success rate is 0.75, p-value < 0.01). An excess of loser to gainer substitutions in the evolution of the omicron line shows that the mutational spectrum has a predictive strength even on the amino-acid level.

### 5. Mutational spectra have long-term amino-acid consequences for all single-stranded positive RNA viruses

The codon usage of the SARS-CoV-2 reference genome was affected by the mutational bias (Fig 3A) and thus the codon usage of other viruses may shed light on their mutational processes. To test it, we analysed genome-wide codon usage of all completely sequenced single-stranded positive RNA viruses affecting humans (N = 149, including separate segments for segmented viruses). We estimated the nucleotide composition in SYN4F sites of these viruses and performed the principal component analysis based on the frequency of A, U, G and C in SYN4F of these 149 viruses. We observed that all coronaviridae form a separate cluster, characterised by the increased fraction of U (Fig 5A, segmented viruses like influenza are presented on this plot with several segments).

**Fig. 5.**
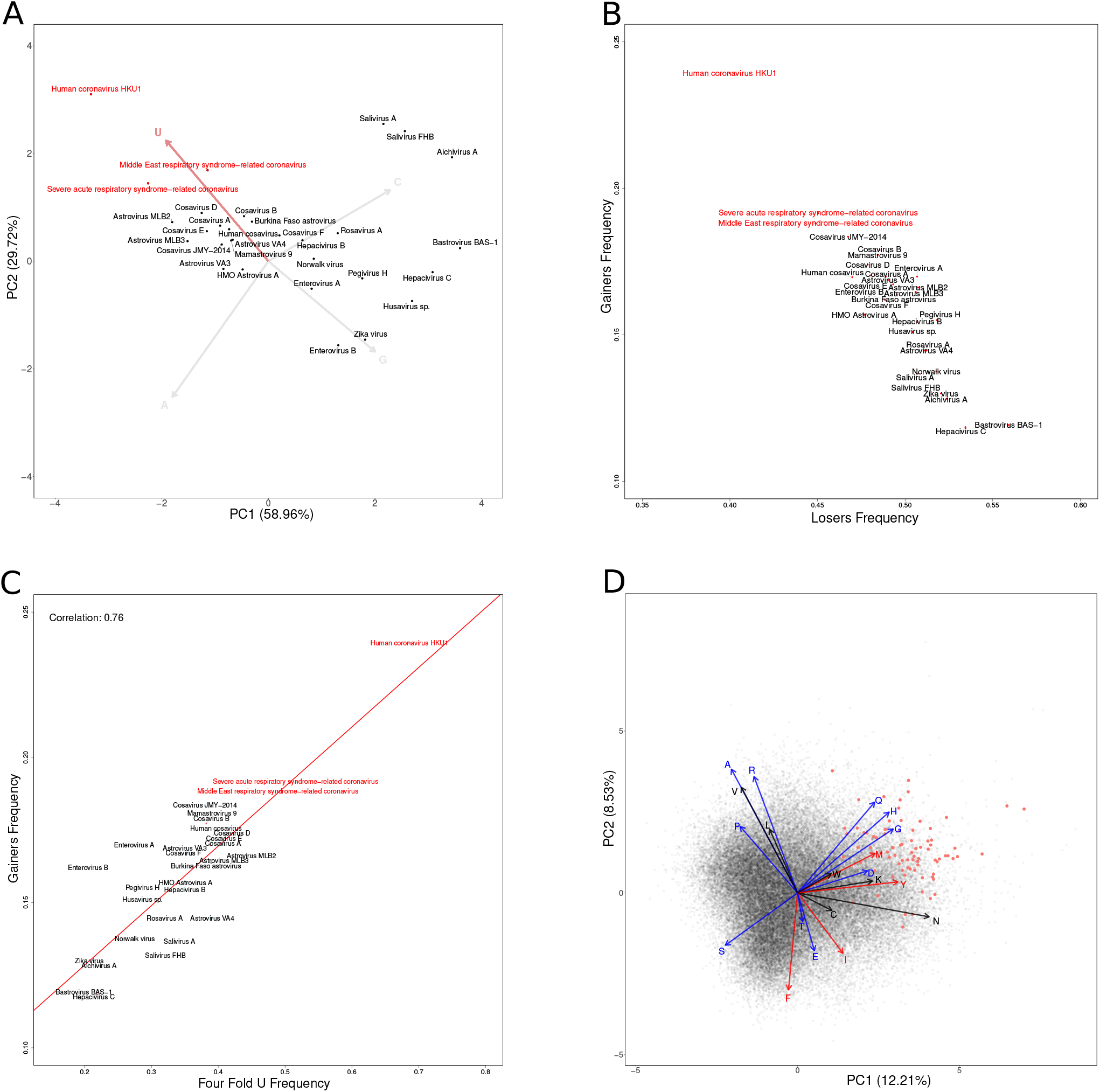
Mutational spectra have long-term amino-acid consequences for all single-stranded positive RNA viruses. All viruses belonging to Coronaviridae family are marked in red colour. 5A Principal component analysis of the neutral (SYN4F) nucleotide content of all completely-sequenced human single-stranded positive RNA viruses. 5B Whole-genome frequencies of gainers and losers among all completely-sequenced human single-stranded positive RNA viruses. 5C Positive correlation of gainers frequency with neutral U frequency (SYN4F) in all completely-sequenced human single-stranded positive RNA viruses. 5D Frequencies of gainers and losers in RNA-dependent RNA polymerase gene of 60793 single-stranded positive RNA viruses. Coronaviridae as compared to other Nidovirales and all +ssRNA viruses are characterized by an excess of gainers and a deficit of losers. 5E Principal component analysis of amino-acid content of RNA-dependent RNA polymerase gene from 60793 single-stranded positive RNA viruses (species-like operational taxonomic units from serratus database). Arrows show a loading of each amino-acid on the first and the second principal components: red marks gainers, blue marks losers, black marks intermediate and neutral amino-acids including leucine (since without the codon structure we can’t distinguish Leu(CUV) from Leu(UUX)).

If mutational bias affects amino acid composition (Fig 4), we can use also comparative species data on amino acid content to proxy the mutational process. We expect that an excess of U in neutral positions (in coronaviridae, Fig 5A) will be linked with an excess of U in non-neutral sites (i.e. excess of gainers in coronaviridae). Oppositely, U deficiency in neutral positions of other species (for example, in Bastrovirus BAS−1, Hepacivirus C, Zika virus and others, see Fig 5A) will be associated with U deficiency in a non-neutral position of these species (excess of losers). To test this, we performed two analyses. First, we estimated the whole-genome amino acid composition of all human single-stranded positive RNA viruses and derived a fraction of gainers and losers for each species. We observed an expected strong negative correlation between gainers and losers, with coronaviridae characterised by an excess of gainers, while viruses such as Bastrovirus BAS−1, Hepacivirus C, Zika virus (with low frequency of U in neutral positions) - by an excess of losers (Fig 5B). This suggests that the mutational process affects the global amino acid composition of human single-stranded positive RNA viruses. Second, taking into account an excess of U among neutral (four-fold degenerated) sites of coronaviridae (Fig 5A) and an excess of gainers among all amino-acids of coronaviridae (Fig 5B) we expect to observe a positive correlation between the frequency of U in neutral positions and the frequency of gainers (which is essentially a frequency of U in not-neutral positions). Indeed, the positive correlation (Fig 5C) confirmed our hypothesis about a link between mutational spectrum and amino-acid composition.

In the analyses above (Fig 5B, Fig 5C) we used each viral genome’s whole amino-acid composition, ignoring potential gene-specificity (different viruses have different sets of genes with different amino-acid composition). To overcome this limitation, we additionally analysed a recent database (30)which focused on the RNA-dependent RNA polymerase (RdRP) gene in more than 10^5 RNA viruses (species-like operational taxonomic units). In the scope of the current project, we focused on all (N = 60793) positive single-stranded RNA viruses (+ssRNA) and estimated an RdRP-specific amino-acid composition of each virus. First, we observed that coronaviridae, compared to other Nidovirales and all +ssRNA viruses, are characterized by an excess of gainers and a deficit of losers (Fig 5D). Second, performing PCA, we observed that both cumulative vectors of all gainers and all losers have high enough projections on pc1 and pc2, suggesting that the mutational spectrum is an essential factor in the molecular evolution of the RdRP gene (Fig 5E). Interestingly, some amino acids, which formally do not support our expectations, can still be driven by mutagenesis. For example, three of four loser amino-acids, showing strong excess in coronaviridae (see four blue arrows with up-right direction Fig 5E): His, Gln and Asp (all except Gly) are in line with the G>A substitution which is the fourth most common substitution in the 12-component mutational spectrum (Fig 1A). This suggests that a more detailed analysis of the effect of the mutational spectrum on amino-acid composition can uncover an even stronger effect of mutagenesis on molecular evolution. Also, despite the observation that Coronaviridae are among the viruses with the highest frequency of gainers, another order - Potatovirales, is even more enriched in gainers. This suggests that a link between life-history traits of different viruses and their specific mutational spectra can be an interesting gap to fill in the future. Altogether, our results align with our hypothesis and show that species-specific mutational spectrum and species-specific amino-acid composition can be tightly linked. Future analyses are needed to shed light on the effect of the mutational spectrum on the amino-acid composition.

## DISCUSSION

Analyzing more than 542768 phylogenetically polarized single-nucleotide substitutions, accumulated during almost two years of the pandemic (from 24.12.2019 to 14.10.2021), we revealed detailed SARS-CoV-2 mutational spectra based on the reconstructed phylogenetic tree. We hypothesised that most analysed mutations (especially from SYN and SYN4F sets) were selectively neutral; thus, the reconstructed mutational spectra reflect the mutagenic process rather than selection. The congruency of the three spectra (ALL - for all mutations, SYN - synonymous substitutions and SYN4F - synonymous mutations in four-fold degenerate sites) with each other (Fig 1, Fig. S1) suggests that, indeed, all of them are mainly shaped by the mutagenesis. It is important to emphasise that in our approach, we count each new mutation only once - at the time of the origin of a given mutation on a given branch of the tree. Even if this mutation was beneficial and led to the strong expansion of the descendant genomes, it will contribute only one event of mutagenesis to the total mutational spectrum. In the case of deleterious variants, a given genome will not propagate and thus does not contribute to our datasets. Thus, our dataset’s total burden of mutations is mainly neutral or effectively neutral. The high similarity of the observed and expected nucleotide compositions (Fig 2C) and nonuniform codon usage of the reference sequence (Fig 3A) suggests that the ancestral viral genome was affected by a very similar mutational process before entering the human host.

In line with other papers (8, 31) we confirmed strong strand-asymmetry and directionality of the 12-component spectra (Fig 1A, 1C), corroborating that an excess of C>U and G>U substitutions is a hallmark of SARS-CoV-2 mutagenesis. In the case of 192-component spectra, our results show slightly different motifs due to our normalisation. According to several authors (8, 31, 32), the most mutagenic context for C>U was ACU, ACA, UCA, UCU and GCU. More detailed analysis showed that normalization on the trinucleotide content of the SARS-CoV-2 genome wasn’t taken into account and thus, their results were biased by the nonuniform background nucleotide content (excess of [AU]C[AU] motifs in the SARS-CoV-2 genome). Our results show that the most C to U substitution occurred in the [A/C/U]CG context. We would like to emphasise the importance of the normalisation procedure for future works: it significantly facilitates a comparison of mutational spectra obtained from different organisms with various nucleotide content. Historically, the main progress in the reconstruction and interpretation of the mutational spectra has been made in the field of human cancers (33, 34), where normalisation was not important and, thus, not used since the background always was the human genome. Normalisation has become increasingly important in the reconstruction of mutational spectra for different organisms. It will facilitate the comparison of the mutational spectra and their deconvolution into signatures of different mutagens using large-scale databases such as COSMIC and SIGNAL.

Mutational bias, if stronger than selection, can affect amino acid substitutions (Fig 4A). The first results, in line with the effect of the mutational spectrum on the amino-acid composition of SARS-CoV-2, were suggested by Simmond (31), which we took into account and investigated in more detail. The butterfly scheme (Fig 4A) helps to formalise our expectations and run tests. Analyzing amino acid substitutions reconstructed on the SARS-CoV-2 phylogenetic tree (Fig 4C), comparing amino acid frequencies between early and late genomes (Fig 4B) and comparing Wuhan and omicron genomes we observed that the majority of substitutions indeed follow the predicted mutational trajectory: from losers to gainers.

To check the robustness of our hypothesis additionally, we analysed all SARS-CoV-2 nonsynonymous substitutions from the CoV-GLUE database (https://cov-glue.cvr.gla.ac.uk/). We downloaded the complete dataset with 5352 polymorphic (different from the reference sequence) nonsynonymous variants segregating with a frequency of at least 0.0001. The majority of these variants (N = 2897), were in line with the trajectories from losers to gainers (driven by C>U or G>U substitutions), 2065 were neutral to our hypothesis (driven by neither C>U nor G>U nor U>C nor U>G substitutions) and only 390 were from gainers to losers (driven by U>C or U>G) and thus opposite to our trajectories. Altogether, this analysis confirmed strong excess of loser-to-gainer substitutions (N=2987) as compared to the opposite ones (N=390) during the recent evolution of SARS-CoV-2.

An extension of this logic on all +ssRNA viruses (Fig 5) suggests the long-term importance of the coronaviridae-specific mutational spectrum on the coronaviridae-specific amino-acid composition. Although we can’t eliminate the selective argument, the mutational explanation is the most parsimonious one for our main findings. Thus we propose that viral-specific amino acid content can be associated with the viral-specific mutational spectrum due to the accumulation of slightly-deleterious variants (losers to gainers) during the effectively neutral molecular evolution. It is important to emphasise that the accumulation of slightly-deleterious variants is commonly expected in genomes with low effective population size, such as the mitochondrial genome of mammals (35). Here we show that the same effect is visible in viral species with typically very high but strongly fluctuating, effective population sizes. An expansion of the viral population during the pandemic, associated with an increased mutational burden, can be an additional factor in the relaxed purifying selection (20).

Our results are in line with a classical “direction mutation pressure” hypothesis by Sueoka (17, 18)and the seminal paper describing correlations between the nucleotide and amino-acid content (19). In our project, we provided one additional layer of information - a deeply reconstructed mutational spectrum, which helps to draw a causative scheme: mutational spectrum > nucleotide content > amino-acid content. It adds a strong argument that all observed correlations between nucleotide and amino-acid content are caused by the mutational bias and corroborates the “direction mutation pressure” hypothesis. RNA viruses are among the fastest evolving creatures on our planet; thus, uncovering a detailed mutational spectrum and consequences of the spectrum on nucleotide, codon and amino acid content is highly important. Recent large-scale mutational datasets allow for uncovering the permissive trajectories of amino-acid substitutions (23)and understanding better the fitness landscape of different proteins (36). Combined with a deep understanding of the mutational spectrum, such studies can predict the global direction of amino-acid changes. This can help to predict better the spectrum of the possible amino acid substitutions, significantly decreasing the dimensionality of expected variants. Understanding the mutational spectrum can help to uncover also evolutionary constrained regions since ‘under all else equal’ they are expected to have an excess of losers and a deficit of gainers. Oppositely, some regions which look constrained (frozen sequence) in reality can be just mutational cold spots (gainers). The proposed “butterfly effect” when the tiny changes in the pattern of mutagenesis can affect the amino-acid content in effectively-neutral regions of different genomes over millions of years can be a very common evolution scenario in species with an asymmetric mutational spectrum, low effective population size and/or high rate of population expansion.

## ACKNOWLEDGMENTS

Computationally heavy reconstruction of the SARS-CoV-2 phylogeny and mutational spectra by KG, BE, and VY was performed on the computational cluster Makarich in the framework of the Russian Science Foundation No. 21-75-20143. All downstream statistical analyses by AV were supported by the Ministry of Science and Higher Education of the Russian Federation (agreement no. 075-02-2022-872). The project was designed by KP in the framework of the federal academic leadership program Priority 2030 at the Immanuel Kant Baltic Federal University.

## Notes

### Competing Interest Statement

The authors have declared no competing interest.

### Summary of Updates

Discussion extended; updated authors list; title and abstract updated.

