## Supplementary Figures for "Direction mutation pressure of SARS-CoV-2 helps to understand the past and predict the future evolution: C>U and G>U biased mutagenesis forces the majority of amino-acid substitutions to be from CG-rich losers to U-rich gainers"

### The butterfly effect: mutational bias of SARS-CoV-2 affects its pattern of molecular evolution on synonymous and nonsynonymous levels

#### SUPPLEMENTARY FIGURES

##### S1. Mutational spectrum of SARS-CoV-2

a. 12-component mutational spectrum of SARS-CoV-2 reconstructed for synonymous sites

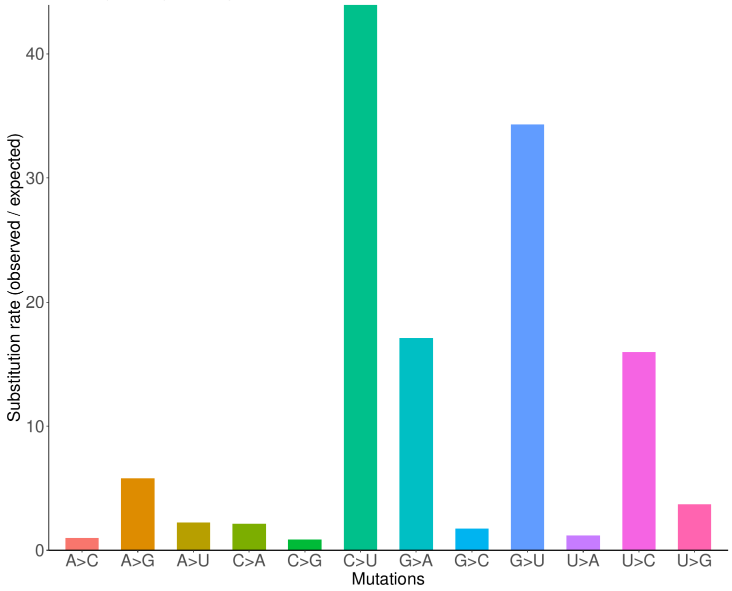


b. 12-component mutational spectrum of SARS-CoV-2 reconstructed for fourfold degenerate synonymous sites

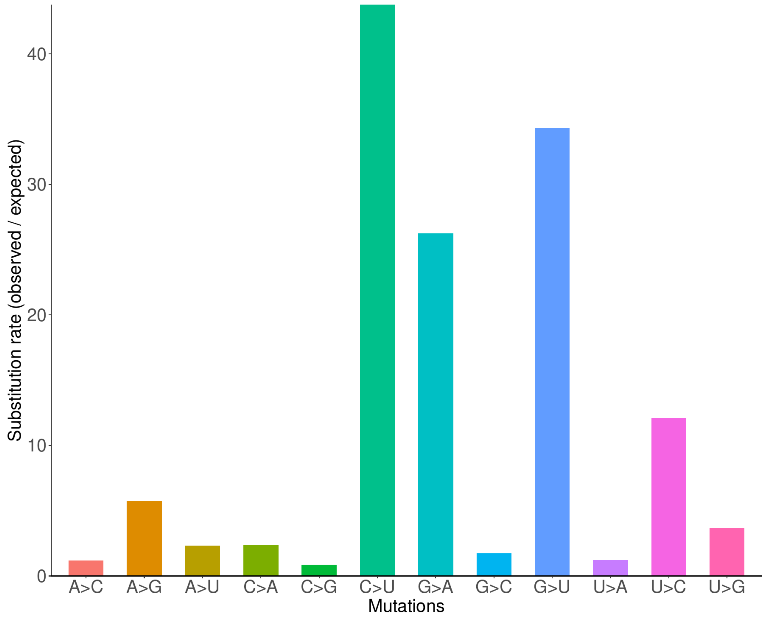

c. A heatmap of the 12-component mutational spectrum of SARS-CoV-2 reconstructed for all sites of each gene separately

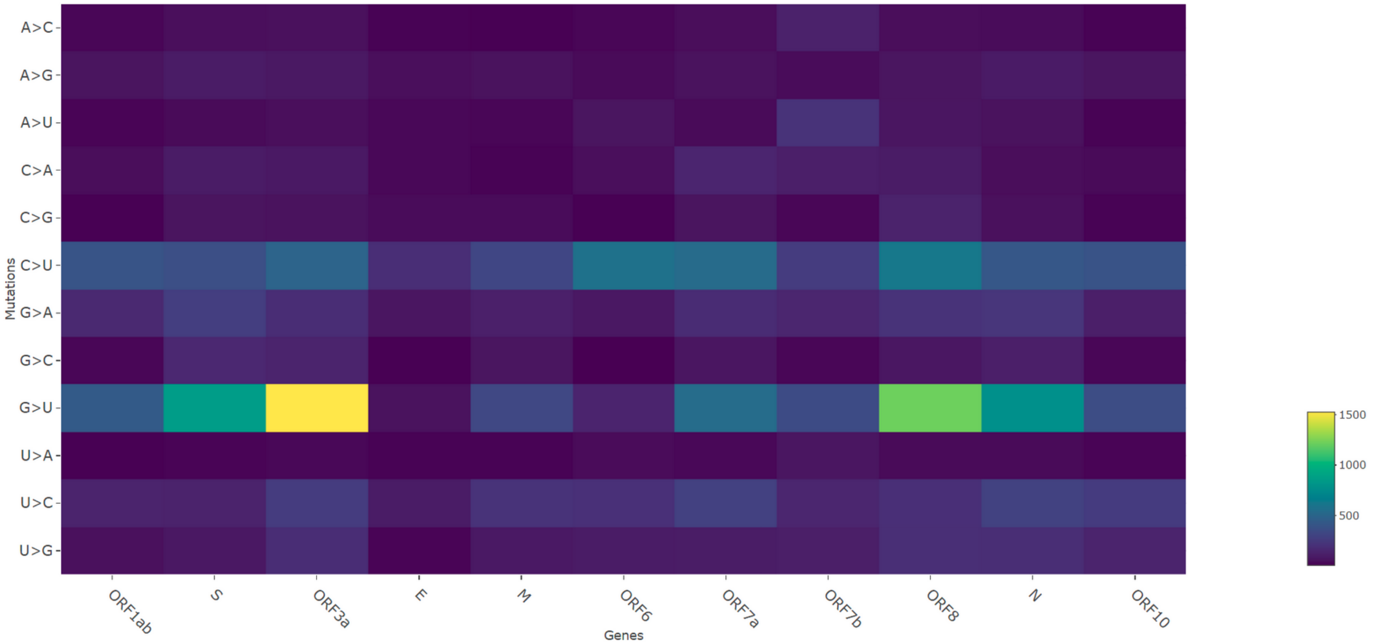

d. Changes in the 12-component mutational spectrum reconstructed for synonymous sites along the pandemic (normalization was performed so that in each time interval, the summa of all mutational frequencies equals one)

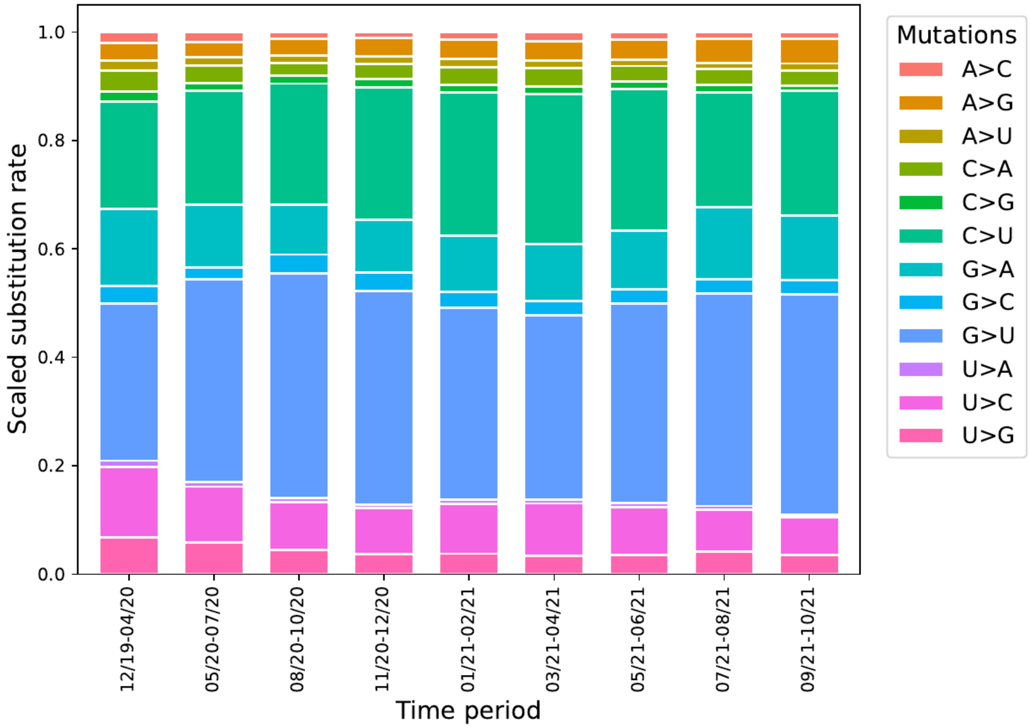


e. Changes in the 12-component mutational spectrum reconstructed for fourfold degenerated sites along the pandemic

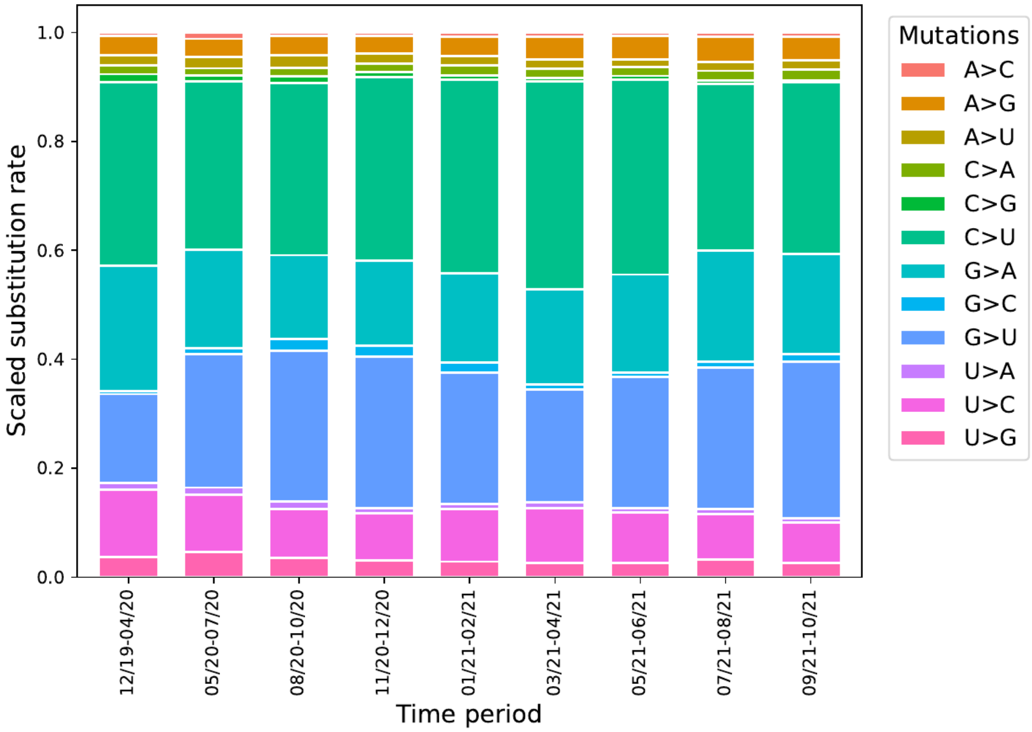

f. 192-component mutational spectrum of SARS-CoV-2 reconstructed for synonymous sites

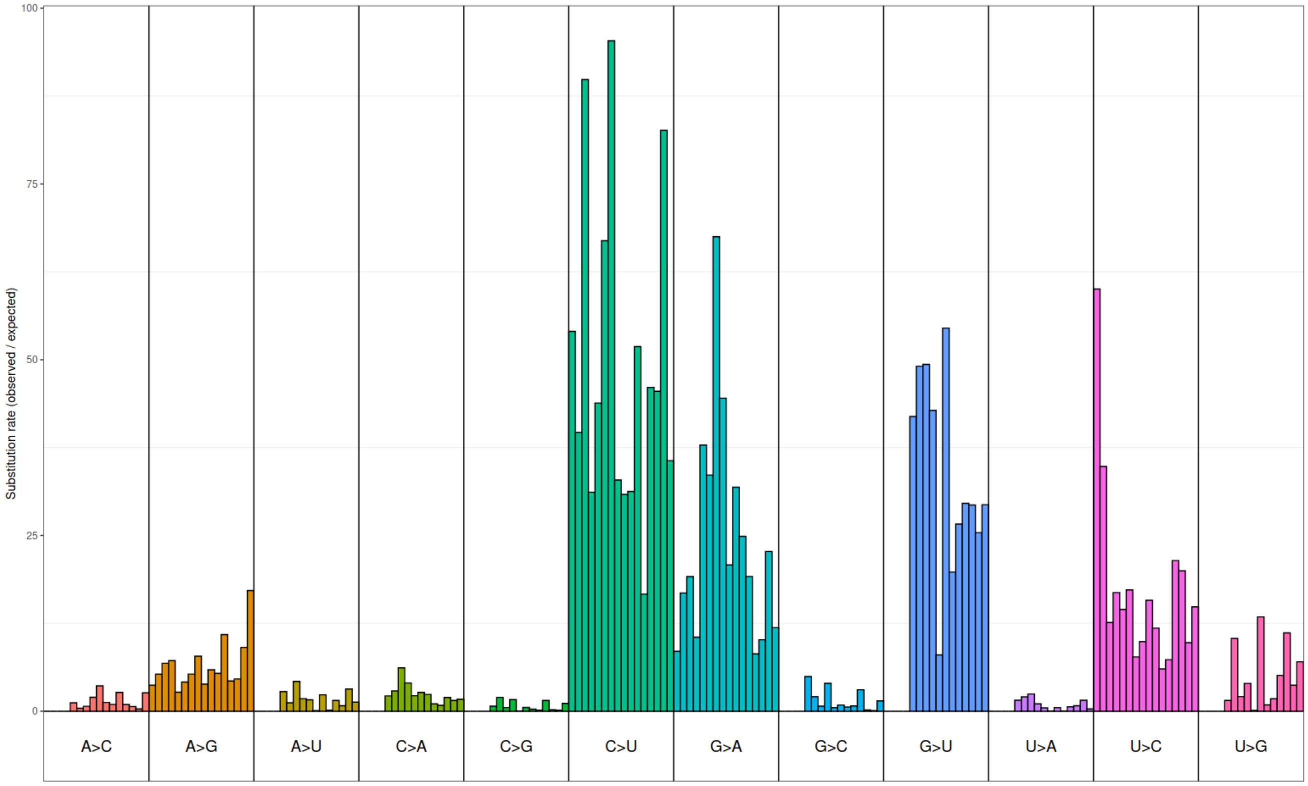


g. 192-component mutational spectrum of SARS-CoV-2 reconstructed for four-fold degenerated synonymous sites

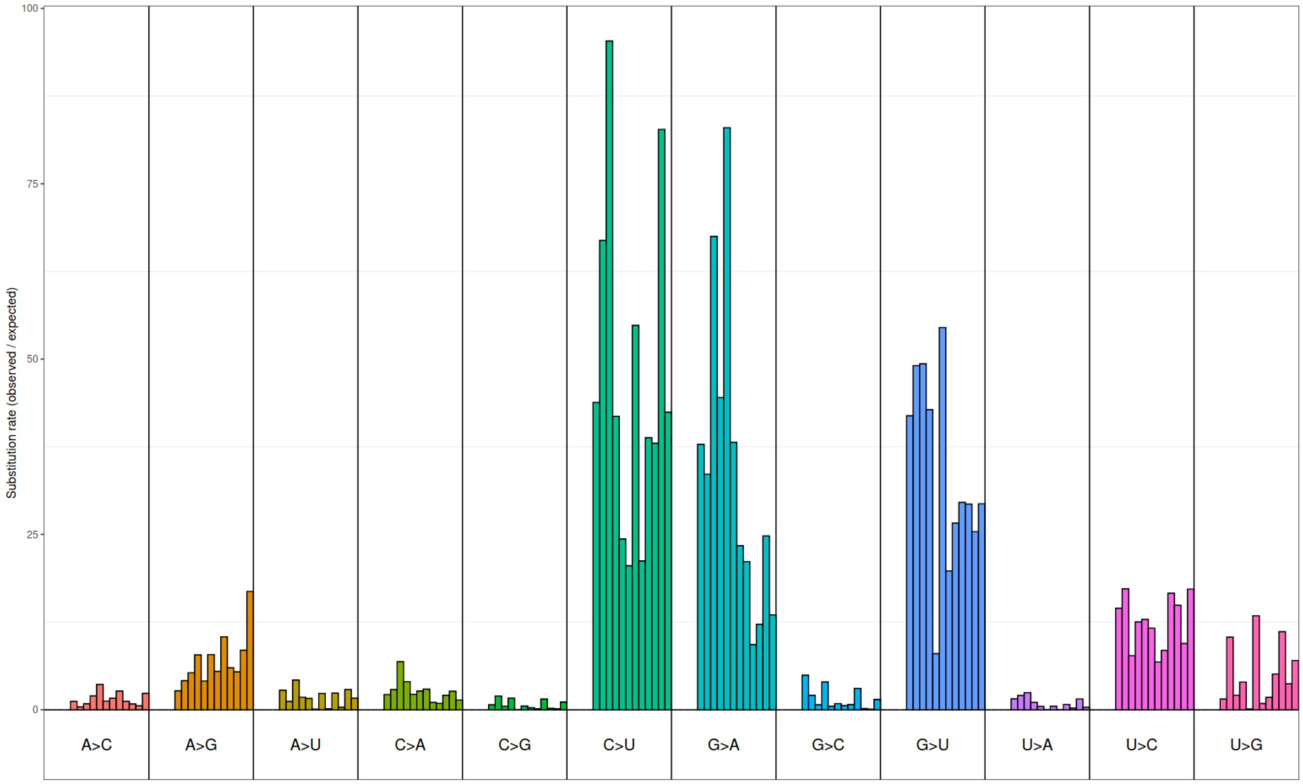

#### S2. Changes in the nucleotide composition of SARS-CoV-2 over pandemic

a. Changes in the nucleotide composition of SARS-CoV-2: four-fold degenerated sites

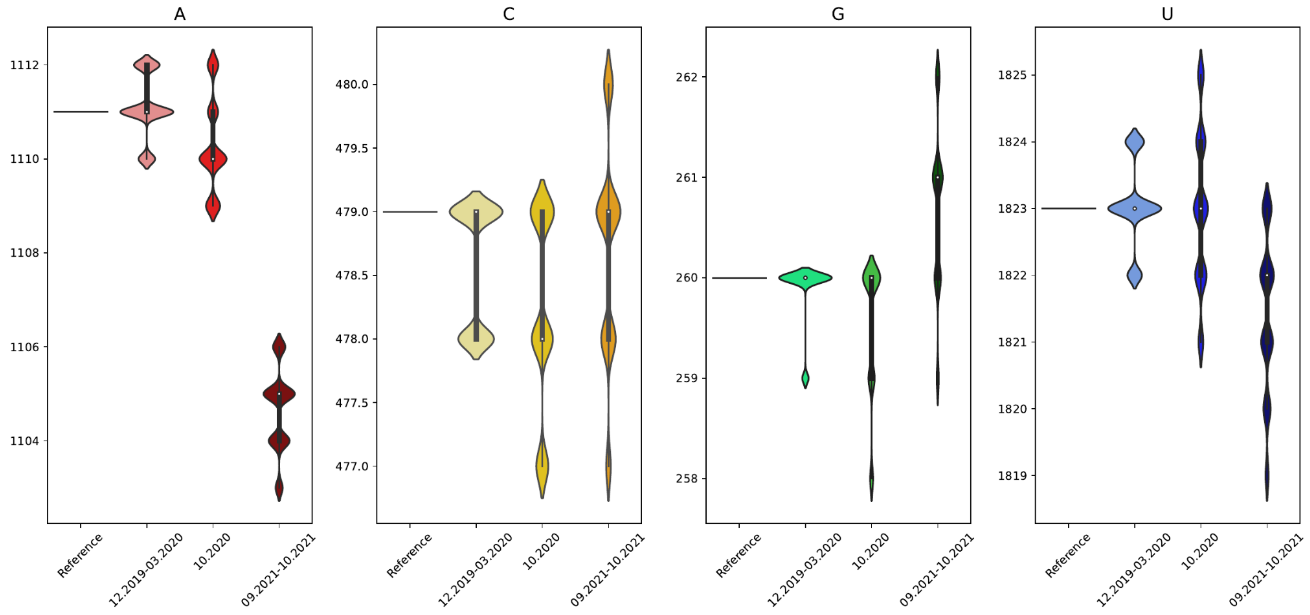


b. Changes in the nucleotide composition of SARS-CoV-2: all sites in XG context

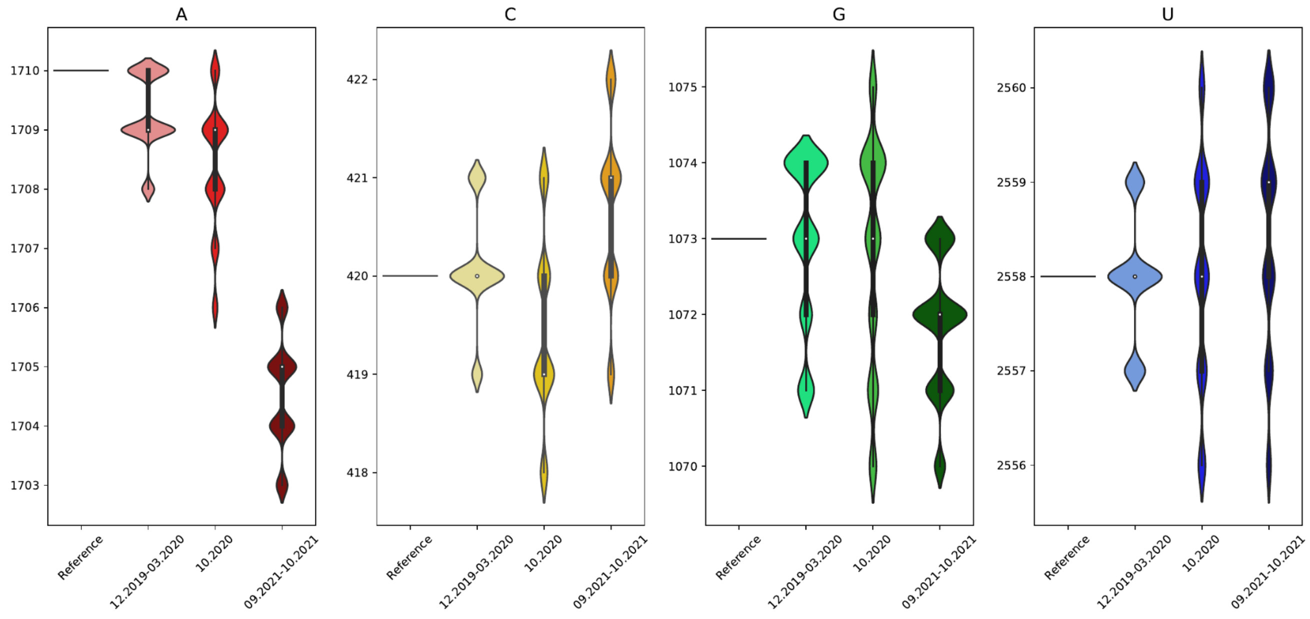

c. Changes in the nucleotide composition of SARS-CoV-2: four-fold degenerate sites in XG context

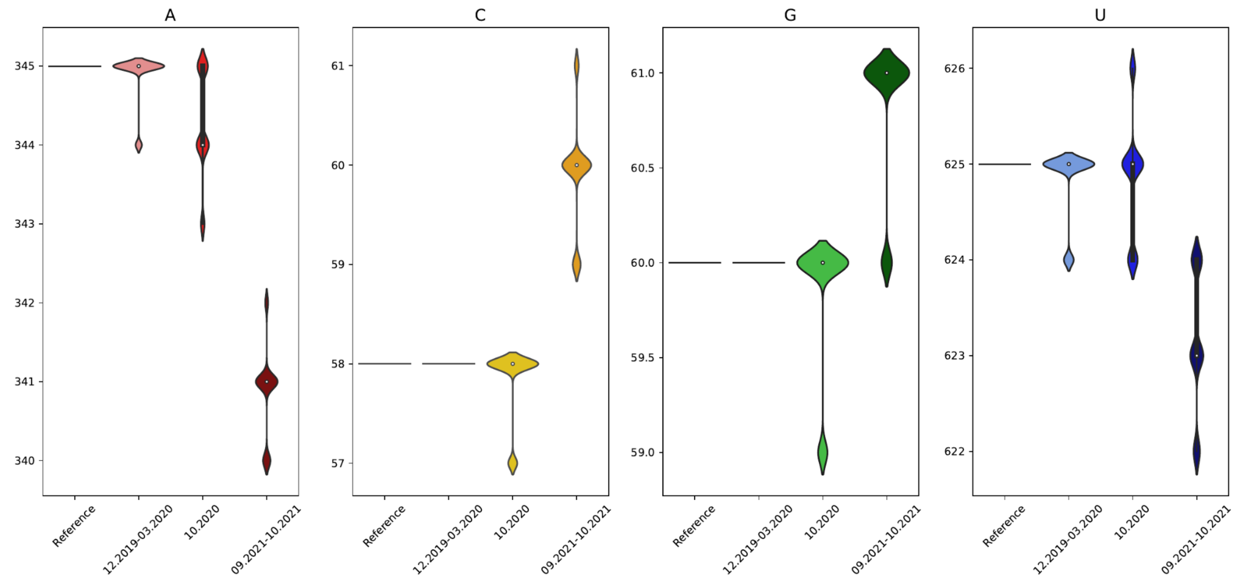

**S3. The codon usage of SARS-CoV-2 is affected by the mutational bias**

The proportion of codons in the reference sequence SARS-CoV-2

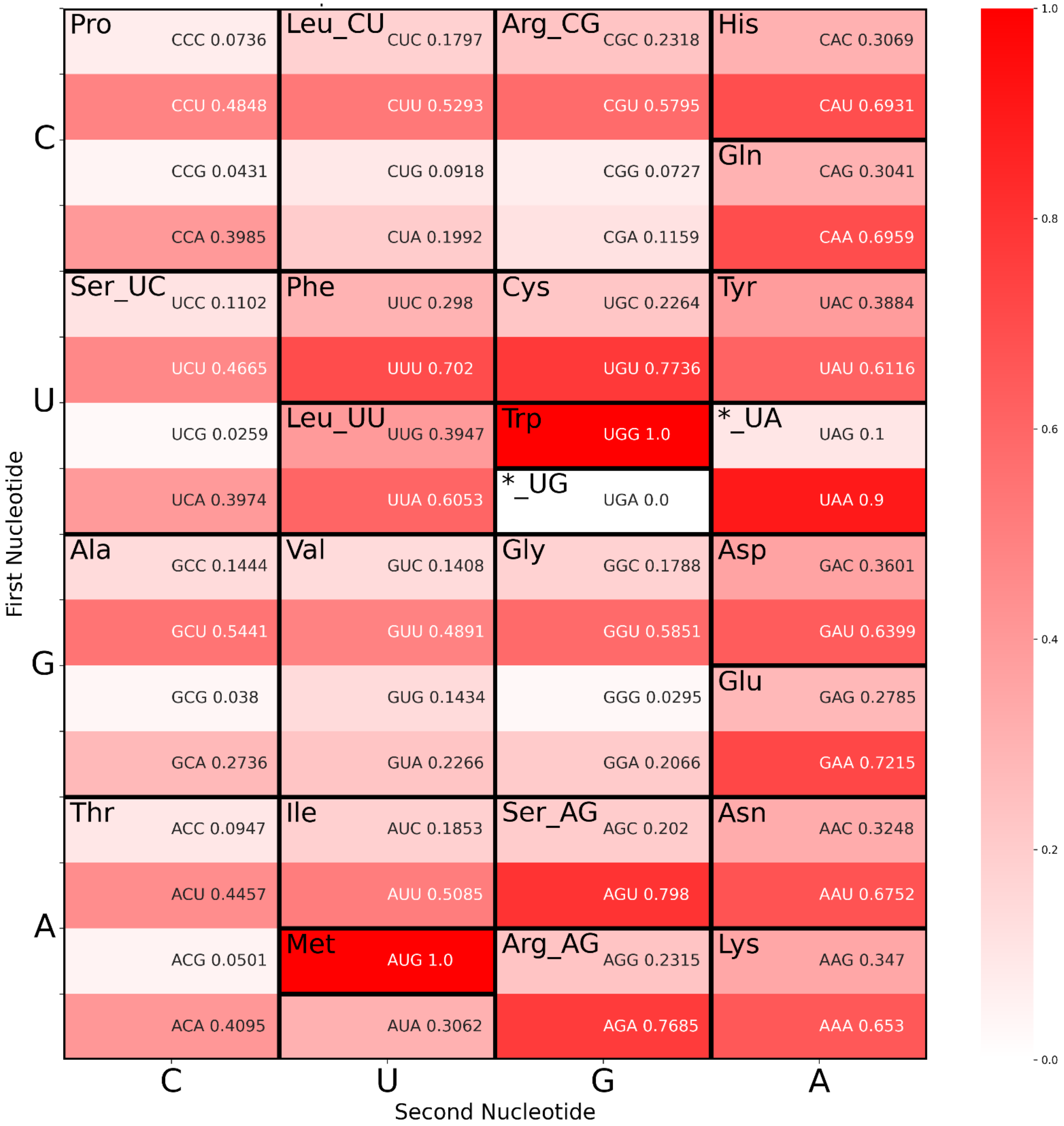

Changes in the codon frequencies during the pandemic

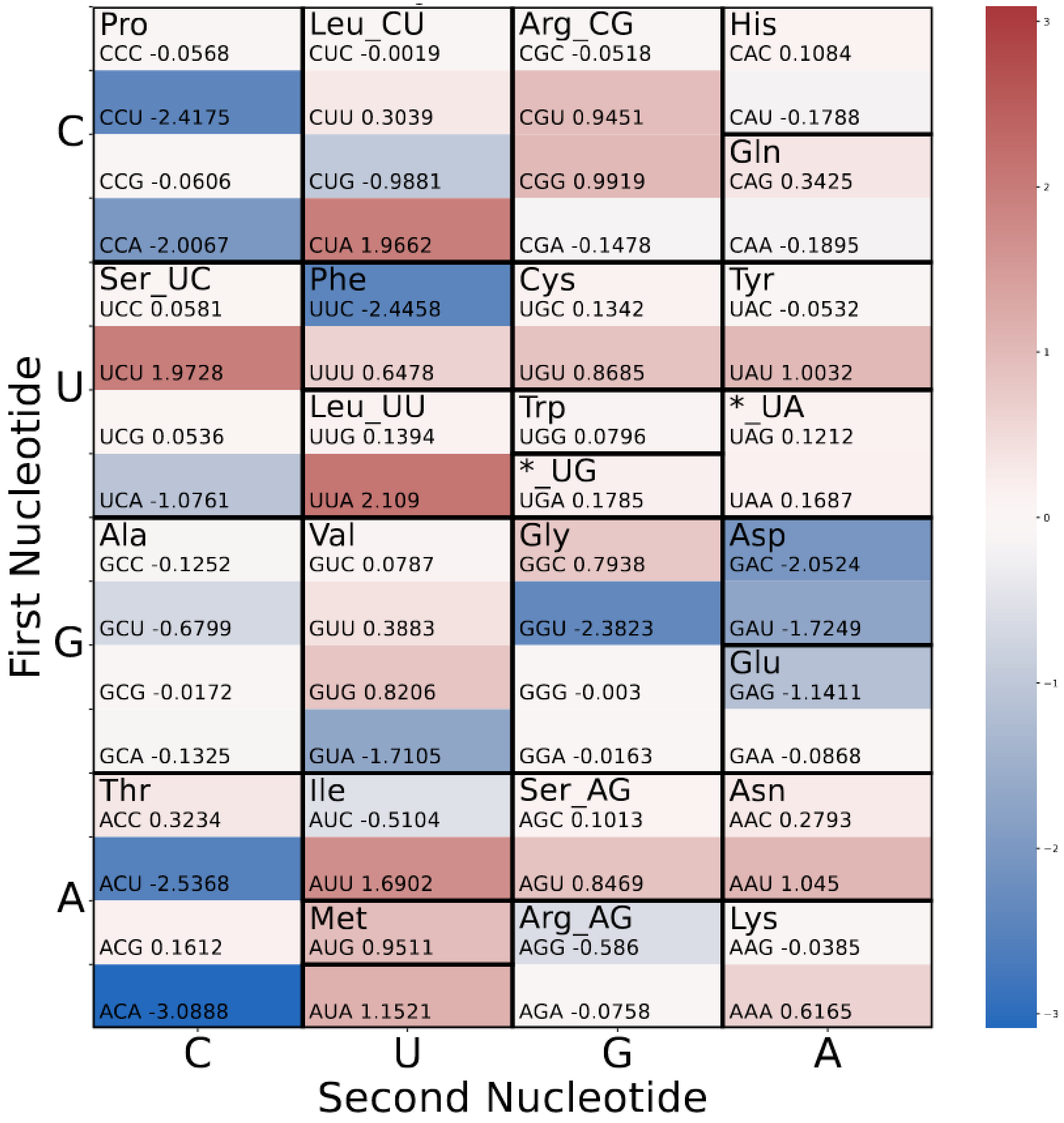

Codon and amino acid usage of all protein-coding genes in the SARS-CoV-2 reference sequence

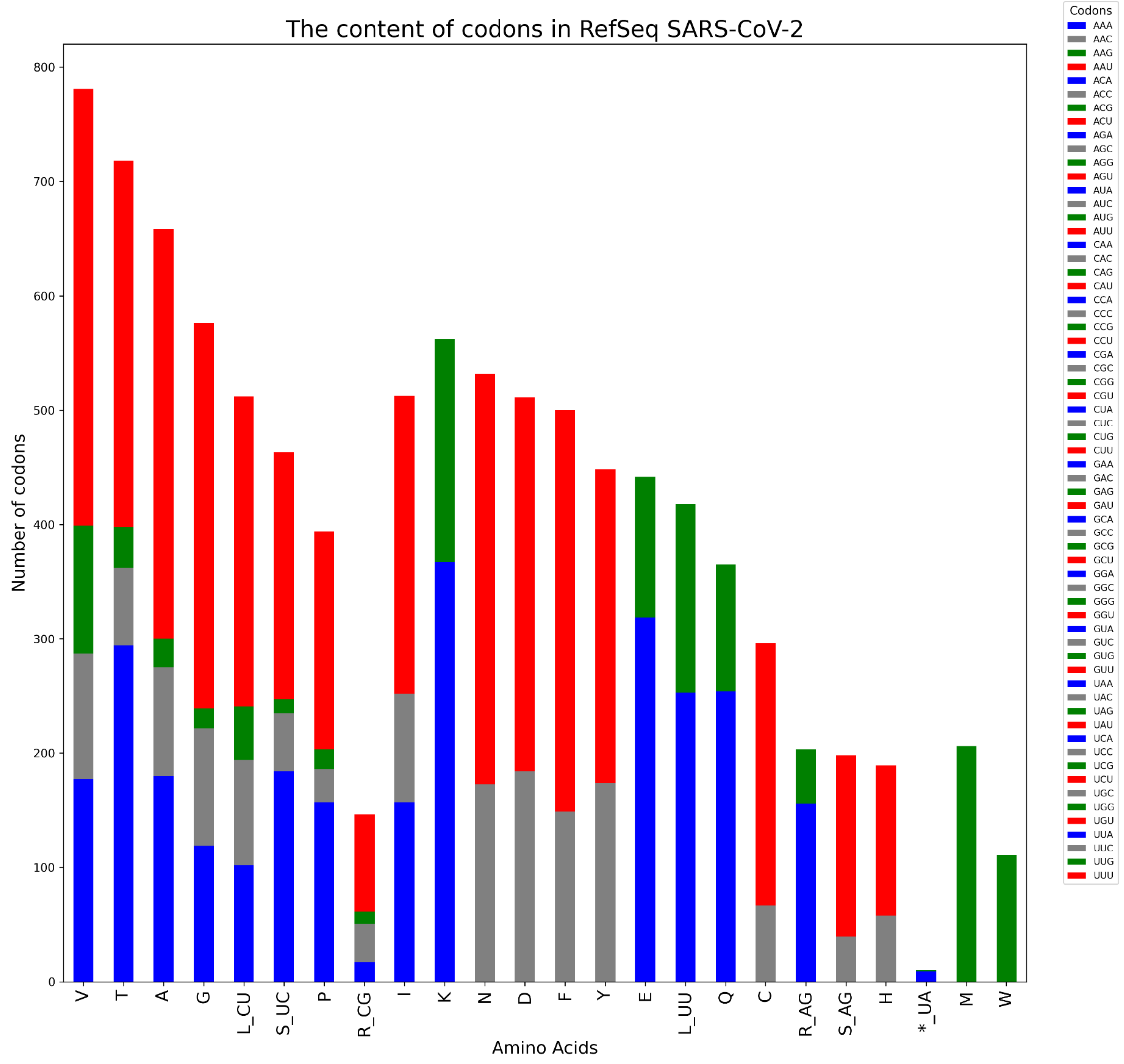

Codon and amino acid usage of ORF1ab in the SARS-CoV-2 reference sequence

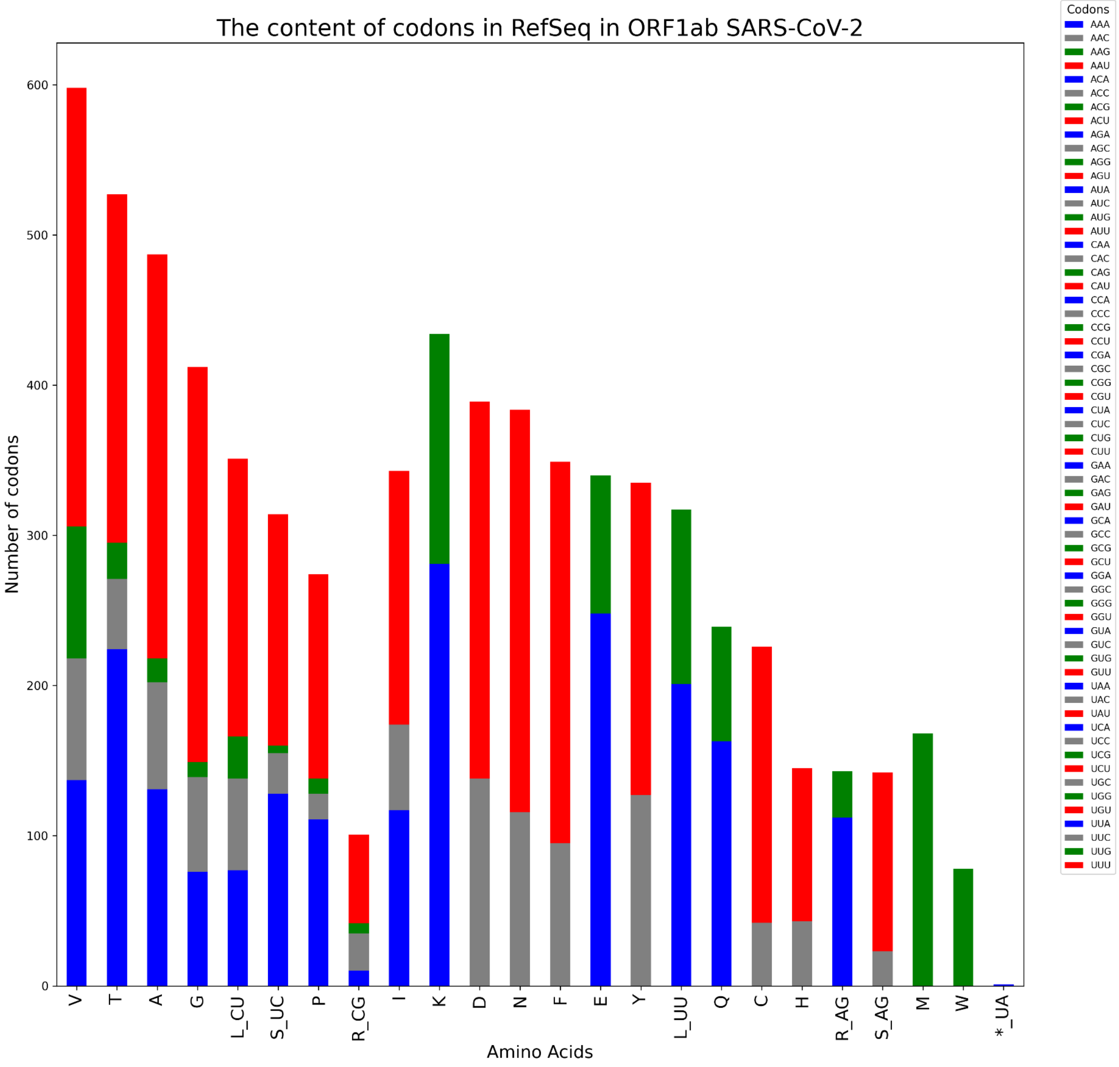

Codon and amino acid usage of all other (all except ORF1ab) proteins in the SARS-CoV-2 reference sequence

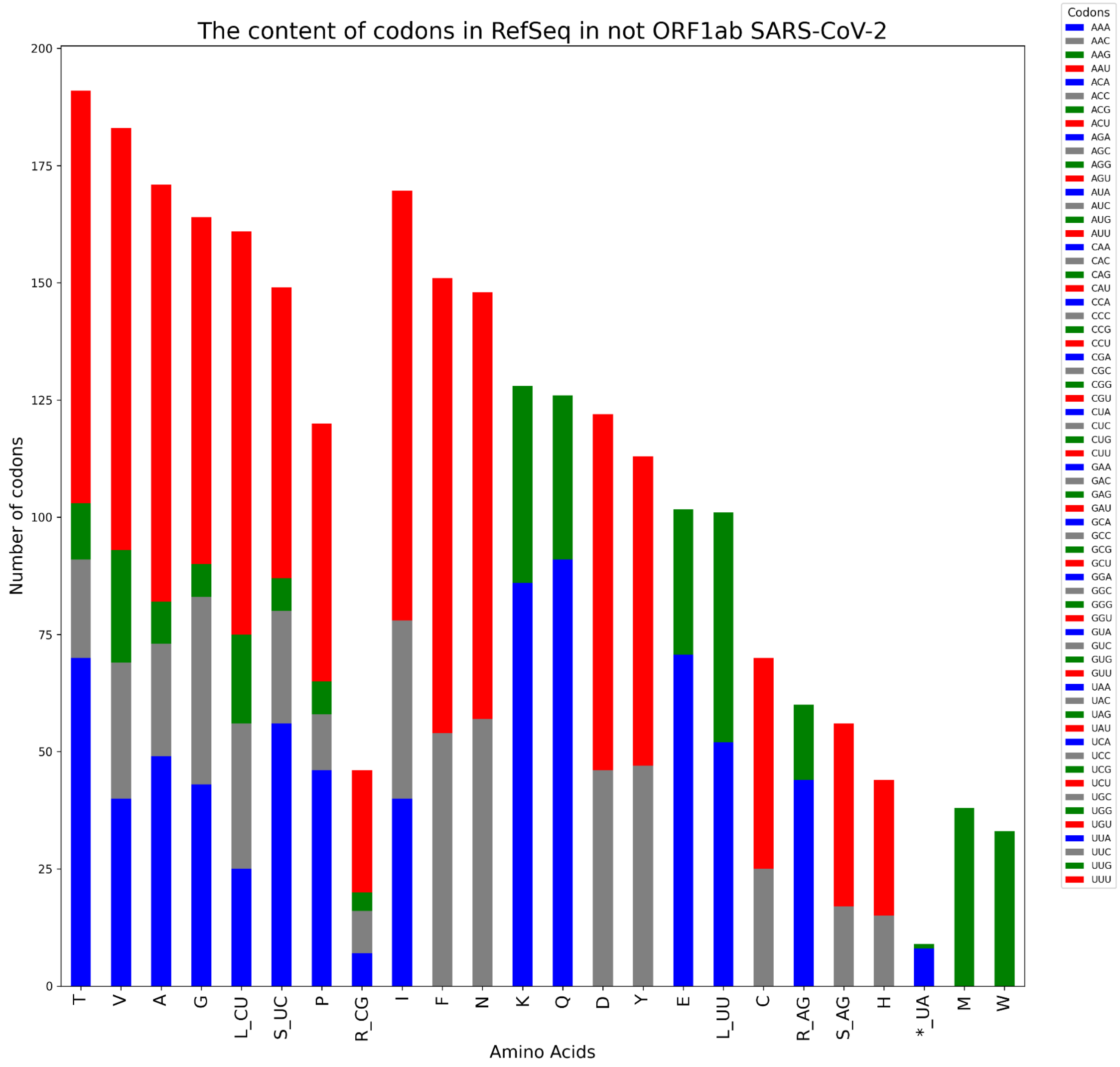

**S4 Amino acid substitutions are affected by the mutational bias**

a. Amino acid composition of the reference genome

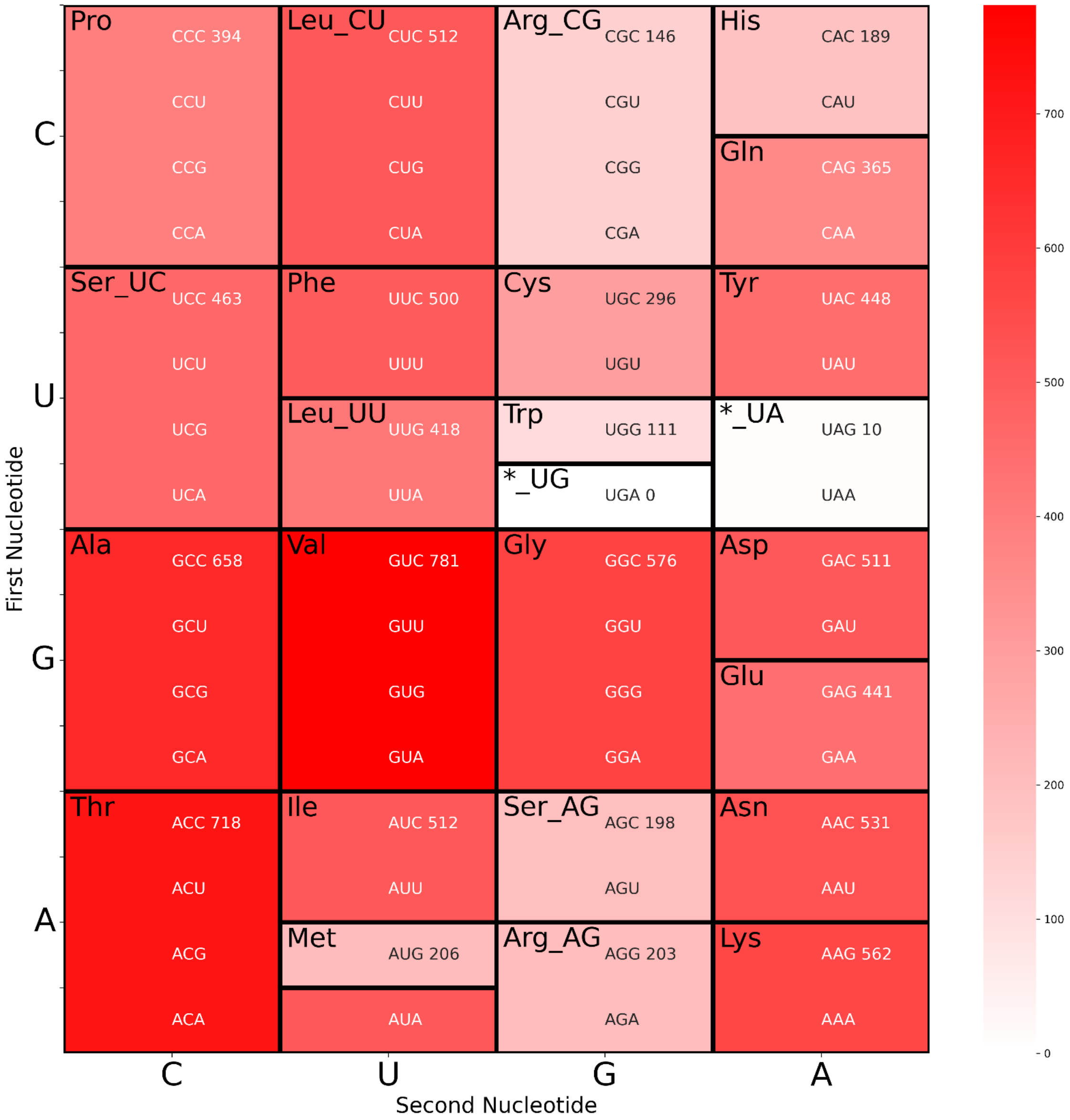

b. Amino-acid substitutions within the ORF1AB gene, reconstructed from the phylogenetic tree, follow the mutational bias

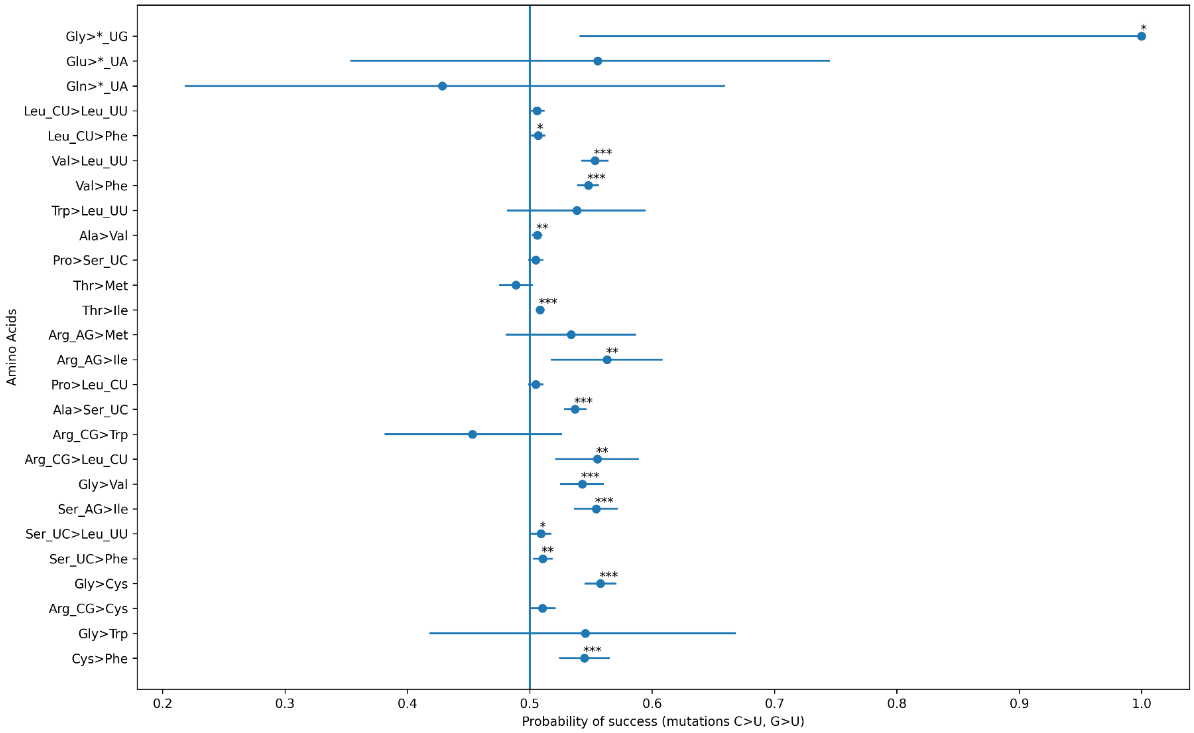


c. Amino-acid substitutions within structural genes (all except ORF1AB gene), reconstructed from the phylogenetic tree, follow the mutational bias

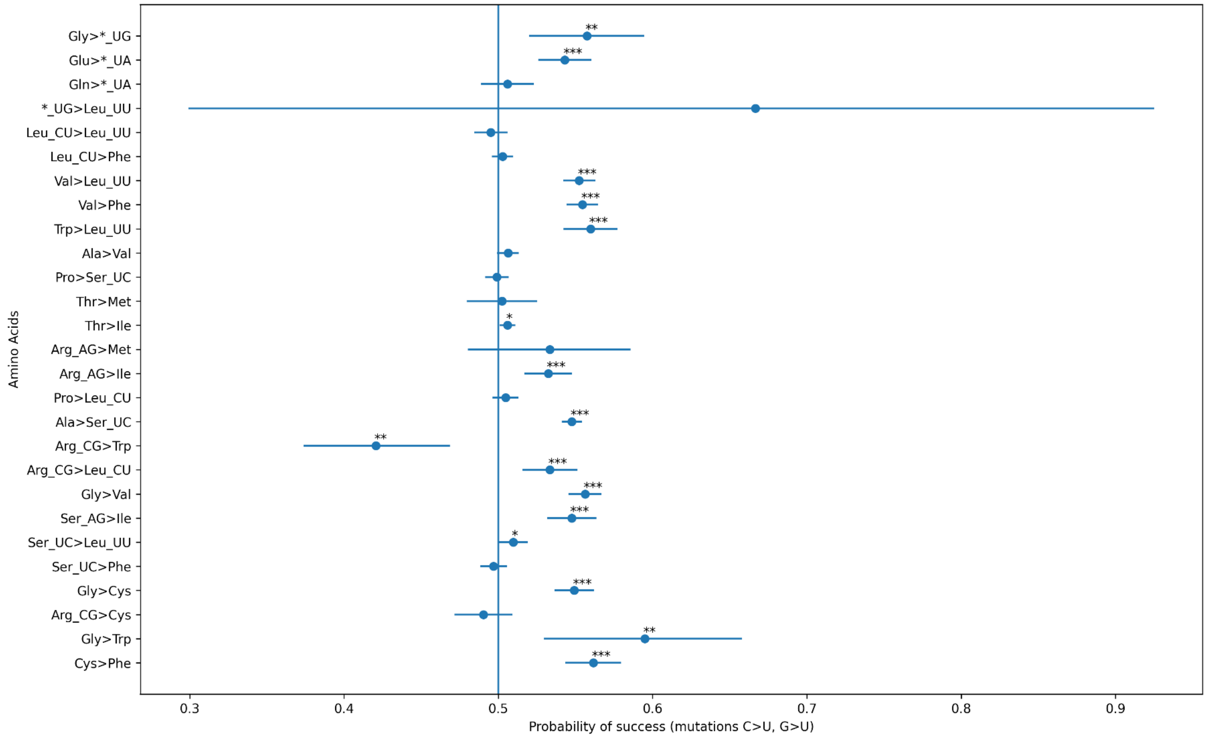
