## Supplementary Tables for "Direction mutation pressure of SARS-CoV-2 helps to understand the past and predict the future evolution: C>U and G>U biased mutagenesis forces the majority of amino-acid substitutions to be from CG-rich losers to U-rich gainers": README.txt

SUPPLEMENTARY TABLES:

Supplemental Table 1. Values of 192-component mutation spectrum based on an ALL set of mutations

Supplemental Table 2. Table with the binomial test for pairs of complementary and reciprocal substitutions for 12 component mutational spectrum of SARS-CoV-2 based on all substitutions

Supplemental Table 3. Table with the binomial test for pairs of complementary and reciprocal substitutions for 12 component mutational spectrum of SARS-CoV-2 based on fourfold substitutions

Supplemental Table 4. Table with the binomial test for pairs of complementary and reciprocal substitutions for 12 component mutational spectrum of SARS-CoV-2 based on synonymous substitutions

Supplemental Table 5. Amino acid substitutions, gainers and losers statistics

Supplemental Table 6. Overall aminoacids substitutions from reference to omicron strain
